# Tissue-specific improvements in CD4^+^ T cell responses after treatment for visceral leishmaniasis

**DOI:** 10.1101/2025.04.13.648617

**Authors:** Jessica A. Engel, Fabian de Labastida Rivera, Benjamin Crawford, Hyun Jae Lee, Jinrui Na, Karshing Chang, Kate H. Gartlan, Luzia Bukali, Teija C.M Frame, Yulin Wang, Ashraful Haque, Christian R. Engwerda

**Affiliations:** QIMR Berghofer Medical Research Institute, Brisbane, QLD 4006, Australia; Department of Microbiology and Immunology, University of Melbourne, located at the Peter Doherty Institute for Infection and Immunity, Parkville, VIC 3000, Australia

## Abstract

Visceral leishmaniasis (VL) is a life-threatening parasitic disease that requires robust CD4^+^ T cell-mediated immunity for parasite control. However, the heterogeneity and transcriptional dynamics of CD4^+^ T cell responses in VL remain poorly defined. In this study, we use a model of experimental VL with tissue-specific immunity and single-cell RNA sequencing to provide a high-resolution assessment of CD4^+^ T cell responses. Our analysis reveals the complexity of CD4^+^ T cell differentiation in VL, identifying distinct Th1 subsets with transcriptional heterogeneity that may reflect functional specialisation. Despite minimal transcriptional differences between splenic and hepatic CD4^+^ T cells, we identified shifts in subset composition, including the emergence of a stem-like CD4^+^ T cell population in the spleen, which was suppressed by the transcription factor Bhlhe40. *Bhlhe40* deficiency skewed CD4^+^ T cell differentiation, impairing Th1 responses while promoting Tr1 cells, resulting in defective parasite control in the liver. Additionally, AmBisome treatment induced a profound transcriptional shift in CD4^+^ T cells, leading to the maintenance of stem-like CD4^+^ T cells in the spleen and the expansion of tissue resident memory-like cells in the liver. These findings uncover key regulatory mechanisms that shape CD4^+^ T cell differentiation in VL and provide insights into how immune-modulatory strategies could enhance long-term immunity.

## Introduction

Visceral leishmaniasis (VL) is a potentially fatal parasitic disease caused by the protozoan parasites *Leishmania donovani* and *L. infantum*. These parasites are transmitted to humans through the bite of infected female sand flies and ultimately infect tissue-resident macrophages in internal organs such as the spleen and liver. VL is endemic in countries in South Asia, East Africa, and South America, and is estimated to cause around 50,000 deaths annually^1^. Despite the availability of effective drugs, VL remains a significant health problem in many parts of the world, particularly in resource-limited settings^2,3,4,5^. There is a need for improved diagnostic tools and more effective treatments, as well as efforts to prevent the spread of the disease through vector control and other measures. However, sustained VL elimination will need an effective vaccine that promotes the development of durable and tissue-specific, anti-parasitic CD4^+^ T cells to prevent outbreaks, especially as case numbers decline and natural immunity wanes as a result of vector control and case-detection measures.

Findings from pre-clinical VL models have shown that following infection, dendritic cells (DCs) capture and present parasite antigens to naïve CD4^+^ T cells in secondary lymphoid tissues, including the spleen^6^. Pattern recognition receptors expressed by DCs are activated by parasite products and trigger IL-12 production that promotes development of pro-inflammatory Tbet^+^ IFNγ^+^ CD4^+^ T helper 1 (Th1) cells^7,8,9^. There is considerable heterogeneity in the molecules Th1 cells express that arises from tissue-specific adaptions for infection control and disease prevention^10^. Regardless, the pro-inflammatory cytokines produced by these Th1 cells are critical for activating phagocytes to kill captured or resident parasites.

The inflammatory products produced by Th1 cells may also damage tissues, and as such, Th1 cells need to be tightly regulated. An important way this occurs is by autologous IL-10 production that inhibits T cell functions and upstream activities of antigen presenting cells (APCs)^11^. IL-10-producing Th1 (type I regulatory T (Tr1)) cell subsets are distinct from Foxp3-expressing regulatory T (Treg) cells that are important for maintaining tissue homeostasis, and comprise a significant proportion of antigen-specific CD4^+^ T cells in VL patients^12^. However, Tr1 cells not only protect tissues, but they may also allow persistent infection by suppressing immune responses via production of anti-inflammatory cytokines, as well as expression of co-inhibitory receptors^13,14,15,16^. Furthermore, they may also restrict responses to vaccination and the effective generation of anti-parasitic immunity^7,15^. Thus, the balance between parasite-specific Th1 and Tr1 cells in infected tissues is an important determinant of disease outcomes.

Given the spleen and liver are important sites of infection and disease in VL patients, much of our knowledge about tissue-specific CD4^+^ T cell differentiation dynamics has come from a pre-clinical model of VL caused by infection of C57BL/6 mice with the human pathogen *L. donovani*^17,18,19^. In this model, the liver is a site of acute, resolving infection with highly effective anti-parasitic CD4^+^ T cells (a balanced Th1 and Tr1 cell response), while the spleen is a site of chronic infection and ineffective CD4^+^ T cells (unbalanced and/or dysregulated Th1 and Tr1 cell response). Thus, the liver responds to infection like infected, asymptomatic individuals (around 90% of *L. donovani*-infected individuals in the Indian sub-continent^3^), while the spleen has a similar disease outcome to that observed in VL patients. The reason for the development of these organ-specific, anti-parasitic CD4^+^ T cell responses is unknown.

In this study, we investigated tissue-specific CD4^+^ T cell responses in C57BL/6 mice following *L. donovani* infection to delineate how CD4^+^ T cell transcriptional landscapes vary across different tissues in acute versus chronic inflammatory environments. By comparing immune responses in distinct tissue sites, we aimed to identify key differences in CD4^+^ T cell dynamics that contribute to protective immunity or immune dysfunction during infection. Additionally, we examined drug-mediated CD4^+^ T cell responses by assessing the impact of anti-parasitic treatment with AmBisome, a liposomal formulation of amphotericin B, which has been linked to the induction of long-lasting immunity in VL patients. This protection is thought to be driven by the maintenance of antigen-experienced CD4^+^ T cells capable of sustaining anti-parasitic immunity^19^. However, how AmBisome treatment influences the preservation or reshaping of CD4^+^ T cell populations, particularly in the context of chronic infection, remains unclear. By distinguishing between tissue-specific and treatment-induced changes in CD4^+^ T cell responses, we aimed to elucidate the mechanisms underpinning durable immunity following infection and therapeutic intervention. Our findings enabled the construction of a high-resolution transcriptomic atlas of CD4^+^ T cell responses in experimental VL, capturing the gene expression landscapes across diverse tissue microenvironments. We identified distinct transcriptional programs that define CD4^+^ T cell subsets in acute and chronic infection, as well as those shaped by anti-parasitic drug treatment. This analysis uncovered molecular signatures distinguishing effector, regulatory, and stem-like CD4^+^ T cell populations, providing a framework for understanding their functional roles. Additionally, we characterised the dynamic transitions in CD4^+^ T cell states during infection resolution, offering deeper insights into the regulatory mechanisms that govern CD4^+^ T cell-mediated immunity in VL.

## Results

### Single cell RNA sequencing reveals a heterogeneous Th1 cell response during experimental VL

To understand how CD4^+^ T cell responses are shaped by acute versus chronic inflammatory environments, we performed transcriptomic profiling of genetically modified T cell receptor transgenic (PEPCK) CD4^+^ T cells in experimental VL^20^. PEPCK is a naturally processed immunodominant antigen derived from *Leishmania* glycosomal phosphoenol-pyruvate carboxykinase (PEPCK_335-351_) conserved in all pathogenic *Leishmania* species^21^. We transferred 10^4^ naïve PEPCK cells from CD45.2 C57BL/6 mice to CD45.1 C57BL/6 mice and infected one day later with *L. donovani*. Using the CD45.2 congenic marker, we isolated transferred PEPCK cells and captured single cell transcriptomes of PEPCK cells from the spleen and liver on day 0, 14, 21,28 and 56 post-infection (p.i.) and performed droplet-based single cell RNA sequencing (scRNAseq) (Figure 1a; Extended Data Figure 1a). In addition, to examine how drug treatment modulates CD4^+^ T cell responses (required for drug-mediated, life-long protection against reinfection^22^), we treated a group of mice with AmBisome, the current recommended VL therapy for patients in India^4^, or vehicle control. In comparison to vehicle control-treated mice, AmBisome treatment led to significant reductions in organ weight and total cell numbers (Extended Data Figure 1b and c), indicating effective parasite clearance and resolution of infection-associated inflammation.

**Figure 1.**
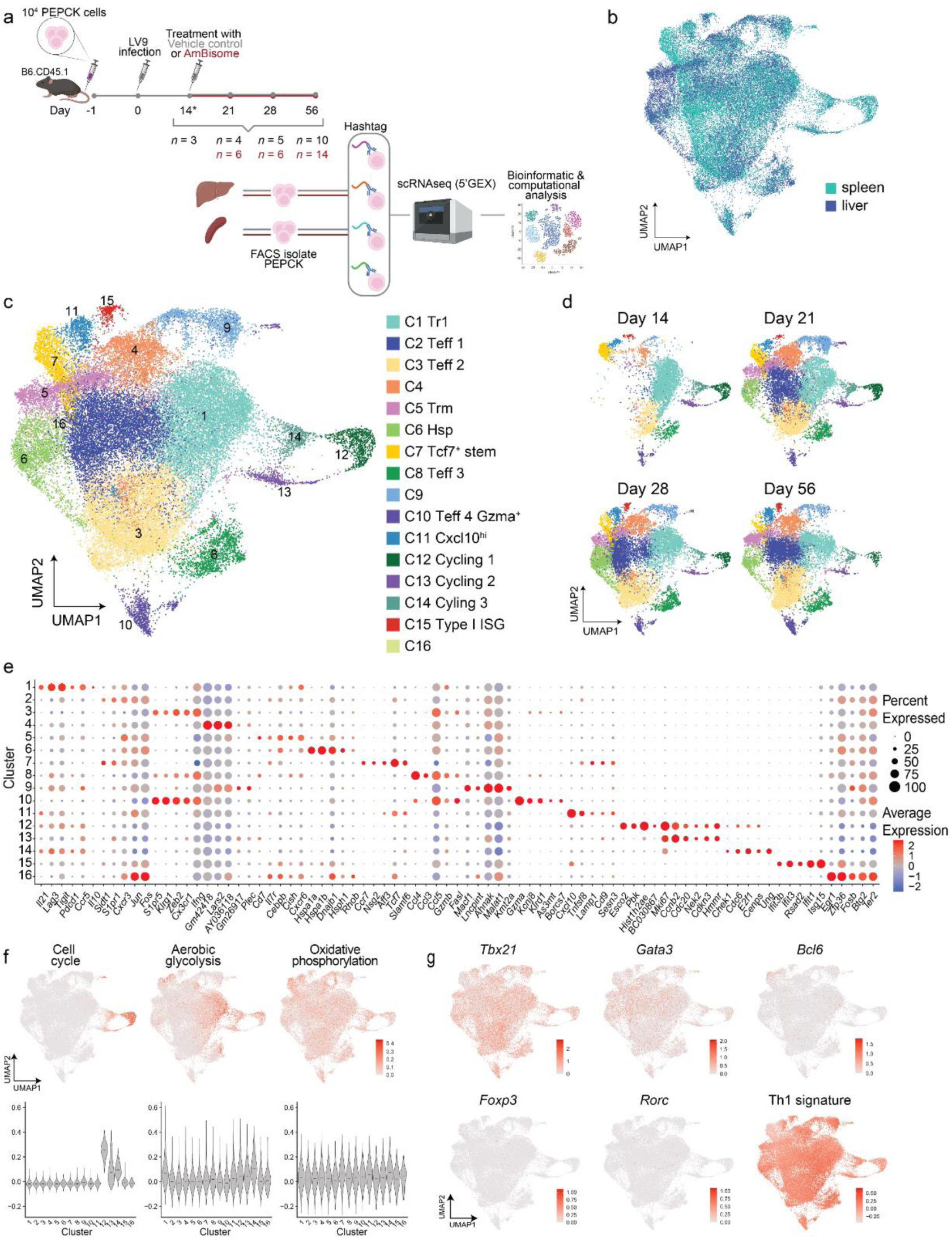
scRNAseq reveals a heterogeneous Th1 cell response during experimental visceral leishmaniasis. **(a)** 10^4^ PEPCK cells (CD45.2^+^) were adoptively transferred to B6.CD45.1^+^ mice, which were then infected with L. donovani the next day. PEPCK cells were isolated from the spleen and liver at various time points post-infection and subjected to scRNAseq. **(b)** UMAP visualisation of 40,667 PEPCK from day 14 – 56 p.i. in experimental VL showing tissue, and **(c)** unsupervised clustering analysis. **(d)** UMAP visualisation showing clusters subsetted by time point p.i.. Colours correspond to clusters in (c). **(e)** Dot plot showing the average expression of curated marker genes for each cluster. **(f)** Cell cycle, aerobic glycoloysis and oxidative phosphorylation gene signature scores visualised on UMAP and quantified in violin plots. Median line shown on violin plots. **(g)** UMAP visualisations showing the expression of T helper cell lineage-defining transcription factors and a Th1 cell gene signature.

To control for batch effects between time points, naïve PEPCK transcriptomes were captured at each time point. Principle component analysis (PCA) revealed that naïve PEPCK transcriptomes from each time point overlapped indicating no batch effects from time point were present in the dataset (Extended Data Figure 2a). After performing quality control checks for naïve and activated genes (*Sell, Ccr7, Cd44, Tbx21*) and due to the transcriptomic distance of naïve PEPCK from all other time points (Extended Data Figure 2b and c), naïve PEPCK were removed from downstream analysis. In addition, to focus on transcriptomic heterogeneity, T cell receptor alpha and T cell receptor beta genes were removed from the dataset prior to further analysis and clustering.

After quality control, 40,667 cells were taken forward for analysis. UMAP visualisation revealed substantial overlap in CD4^+^ T cell transcriptomes between acute (liver) and chronic (spleen) inflammatory environments (Figure 1b), suggesting shared transcriptional programs across tissues. Unsupervised clustering defined 16 clusters (Figure 1c; Extended Data Figure 2d & e), which included 3 main clusters (>10% of total cells; Clusters 1-3), 6 intermediate clusters (3-10% of total cells; Clusters 4-9) and 7 minor clusters (<3% of total cells; Clusters 10-16). All clusters were observed by day 14 p.i. with little evidence of progressive change after day 21 p.i. over the course of infection (Figure 1d), indicating that patterns in CD4^+^ T cell responses are established early during infection and do not undergo significant diversification as the infection progresses. Cell clusters were annotated using cluster marker genes (Figure 1e) and included; effector cells expressing *Cxcr3, Jun* and *Fos* (Cluster 2), and *Klrg1*, *Zeb2*, *S1pr5*, *Ifng* (Cluster 3), and *Ccl3*, *Ccl4* and *Gzmb* (Cluster 8), and *Gzma* (Cluster 10); Tr1-like cells (Cluster 1) expressing *Lag3*, *Tigit*, *Pdcd1* and *Il10*; memory-like cells expressing *Cd7*, *Il7r, Cxcr6* (Cluster 5) and *Tcf7, Ccr7* (Cluster 7); cells expressing heat shock stress pathway genes (Cluster 6) and type I interferon stimulated genes (Cluster 15); proliferative cells (Cluster 12, 13, 14) expressing cell cycle genes (Figure 1f). In addition, PEPCK cells in the proliferative clusters, as well as in Cluster 1, showed increased aerobic glycolysis activity (Figure 1f). Assessment of the expression of lineage marking transcription factors showed that PEPCK cells in all clusters, but not naïve PEPCK (Extended Data Figure 2c), expressed high levels of *Tbx21* and upregulated genes associated with a Th1 cell gene signature (Figure 1g), consistent with a Th1 cell phenotype, as previously reported in experimental VL^16^. These data suggest that during experimental VL, CD4^+^ T cells diversify into various Th1 cell subtypes with distinct gene expression profiles.

### Transcriptional similarity of CD4^+^ T cells in distinct tissues and reprogramming following AmBisome treatment during experimental VL

To investigate the influence of tissue environment and drug treatment on CD4^+^ T cell differentiation, we examined the distribution of PEPCK cells across clusters. Although the heterogeneity of PEPCK cells was similar in most samples, their relative proportions varied depending on time point, tissue, and treatment (Figure 2a & b; Extended Data Figure 3). Overall, the transcriptional profiles of CD4^+^ T cells in the spleen and liver from control-treated mice were similar, with most clusters following similar temporal dynamics across both organs. Notable organ-specific differences in cluster frequencies included Cluster 6, associated with heat shock stress pathways, which was more prevalent in the liver at day 21 p.i.. Additionally, Cluster 7, comprising *Tcf7*-expressing PEPCK cells, peaked in frequency at day 14 p.i. and was largely spleen-specific. Taken together, these data suggest that CD4^+^ T cells in the spleen and liver exhibit largely comparable transcriptional programs throughout infection, indicating that factors beyond T cell intrinsic transcriptional differences, such as the tissue microenvironment, likely drive organ-specific infection outcomes.

**Figure 2.**
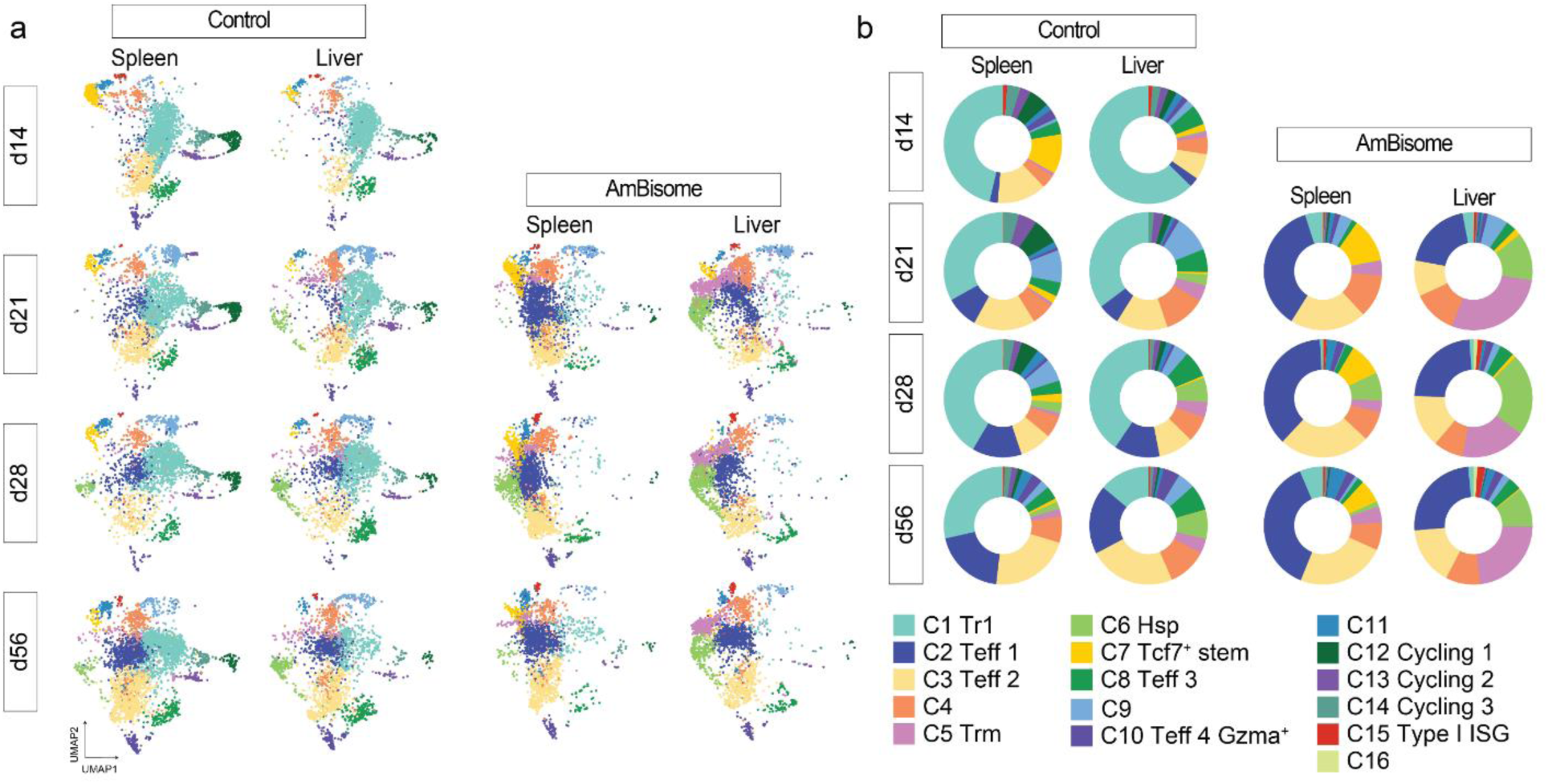
Tissue-specific CD4^+^ T cell responses in experimental VL. **(a)** UMAP visualisation showing PEPCK cells at day 14-56 p.i. in experimental VL subsetted by time point, organ and treatment. **(b)** Donut plots show the proportion of PEPCK cells from each sample per cluster. Colours correspond to clusters in (a).

We next investigated the impact of AmBisome treatment on CD4^+^ T cell transcriptional profiles and their developmental trajectories. Strikingly, AmBisome treatment induced substantial shifts in cluster frequencies, suggesting a pronounced impact on CD4^+^ T cell dynamics. Most notably, the proportion of Tr1 cell PEPCK cells (Cluster 1) was dramatically reduced following treatment, while effector PEPCK cell populations (Clusters 2 and 3) expanded in both the spleen and liver. Additionally, Cluster 7, a *Tcf7^+^* spleen-associated population, was maintained at a higher frequency beyond day 14 p.i. in AmBisome-treated mice, instead of declining, as occurred in untreated controls. In the liver, Cluster 5, which exhibited tissue resident memory (Trm)-like characteristics, expanded significantly following drug treatment. Overall, while some tissue-specific differences were observed in untreated mice, the most striking changes were driven by drug treatment. AmBisome not only sustained the presence of splenic *Tcf7*-expressing cells throughout infection but also promoted the development of Trm-like PEPCK cells in the liver, highlighting its capacity to shape distinct CD4^+^ T cell subsets across tissues.

These transcriptomic findings were validated at the protein level through flow cytometry analysis of PEPCK cells isolated from the spleen and liver of *L. donovani* infected mice at days 14, 28, and 56 p.i., including a cohort treated with AmBisome at day 14 p.i. (Extended Data Figure 4a and b). Flow cytometry revealed substantial heterogeneity in surface marker expression among PEPCK cells, mirroring the diversity observed in transcriptional profiling (Extended Data Figure 4c and d). All PEPCK cells expressed Tbet and CD44, confirming an activated Th1 cell phenotype. TCF1 expression was observed to be the highest in the spleen at day 14 p.i., and was sustained at later time points following AmBisome treatment, compared to in control-treated mice. AmBisome treatment also resulted in reduced PD1 expression in both the spleen and liver, suggesting a potential alleviation of T cell exhaustion. Additionally, AmBisome treatment led to upregulation of CXCR6 and CD69 expression in the liver, consistent with an expansion of tissue-resident cells, while CD127 expression increased in both the spleen and liver, indicative of enhanced survival potential.

To further define these populations, unsupervised clustering of PEPCK cell FACS data identified 18 distinct clusters (Extended Data Figure 4e and f). These included seven effector clusters, two memory-like clusters (tissue-resident memory [Trm] and central memory [Tcm]), and nine TCF1^+^ clusters distinguished by differential expression of CCR7, CD127, CXCR6, PD1, and KLRG1. Consistent with transcriptional data, the frequency of these clusters varied by tissue, time point, and treatment (Extended Data Figure 4g). Overall, these findings confirm the heterogeneity of Th1 cells at the protein level and demonstrate that tissue environment and drug treatment drive distinct Th1 cell subset frequencies within the CD4^+^ T cell response during experimental VL.

### Polyclonal CD4^+^ T cell responses resemble PEPCK cell responses at the transcriptomic level

The diversity of the T cell receptor (TCR) repertoire has been shown to influence CD4^+^ T cell responses by affecting clonal selection, signal strength, and subsequent differentiation into distinct effector and memory subsets^23,24^. In experimental VL, we observed diversity within Th1 cells, but limited transcriptomic differences between CD4^+^ T cells from the spleen and liver. To determine whether these patterns are influenced by lack of T cell receptor (TCR) diversity, we investigated whether T cell clonality drives effector fate choices or tissue localisation, and whether the lack of transcriptomic variation reflects a limited repertoire of TCR diversity. We used scRNAseq to compare the transcriptomes of PEPCK cells (monoclonal) and polyclonal *Leishmania*-specific CD4^+^ T cells during experimental VL. Congenic wild-type mice were infected with *L. donovani* and polyclonal *Leishmania*-specific CD4^+^ T cells were isolated at day 14 p.i. from the spleen and liver by staining with a *Leishmania*-specific MHC-class II I-A^b^ tetramer (I-A^b^ PEPCK_335-351_)^25,26^, and cells were then sorted. This time point is the peak of acute infection in the liver and marks the establishment of infection in the spleen in experimental VL^19^. To minimise non-specific binding to the I-A^b^ PEPCK_335-351_ tetramer, we co-stained with a negative control MHC-class II I-A^b^ tetramer loaded with a non-specific human CLIP peptide (I-A^b^ CLIP_87-101_) (Extended Data Figure 5a). Activation of isolated I-A^b^ PEPCK_335-351_^+^ cells was confirmed by co-expression of CD49b and CD11a (Extended Data Figure 5b) and the transcriptomes were then assessed using droplet-based scRNAseq coupled with TCR V(D)J sequencing using the 10x Chromium platform. Due to constraints in isolating sufficient cell numbers, cells from multiple mice were pooled for scRNAseq as described in the methods. This resulted in single cell transcriptomic data for 5,348 I-A^b^ PEPCK_335-351_^+^ cells (2,935 cells from the spleen and 2,413 cells from the liver) after excluding poor quality single cell transcriptomes (Extended Data Figure 5c), which were then merged with scRNAseq data from splenic and hepatic PEPCK cells at day 14 p.i (from Figure 1).

PEPCK and I-A^b^ PEPCK_335-351_^+^ cells from the spleen and liver showed significant transcriptomic overlap when visualised by UMAP (Figure 3a). Clustering of PEPCK and I-A^b^ PEPCK_335-351_^+^ cells at day 14 p.i. defined 17 clusters (Figure 3b), including 4 main clusters (Clusters 1-4), 6 intermediate clusters (Clusters 5-10) and 7 minor clusters (Clusters 11-17). As defined using cluster marker genes, similar CD4^+^ T cell subsets were identified as before for the PEPCK cell longitudinal data. This included Tr1 cells (Cluster 2) expressing high levels of *Lag3* and *Il10*; effector cells (Clusters 1, 4, 7, 15) expressing effector genes *S100a4*, *Ccl5*, *Nkg7*, *Gzma*; proliferative cells (Clusters 6, 9, 12); memory cells expressing high levels of *Tcf7* and *Ccr7* (Cluster 4). PEPCK and I-A^b^ PEPCK_335-351_^+^ cells from the spleen and liver were broadly distributed across all clusters, excluding Clusters 13, 15, and 17 (Figure 3c). Clusters 13 and 17 were predominantly composed of I-A^b^ PEPCK_335-351_^+^ cells, with Cluster 13 enriched for cells expressing heat shock stress pathway genes and Cluster 17 representing a minor population of only 36 cells. A similar heat shock stress-associated cluster was also identified in the longitudinal PEPCK cell dataset (Figure 1; Cluster 6), but it was only identified from day 21 onwards, suggesting that differences in infection kinetics may contribute to this variation and that this signature is not specific to I-A^b^ PEPCK_335-351_^+^ cells. In contrast, Cluster 15 was predominantly composed of PEPCK cells, suggesting that clonotype may influence the likelihood of CD4^+^ T cells differentiating into *Gzma* expressing effector cells. However, further experimentation is required to confirm this.

**Figure 3.**
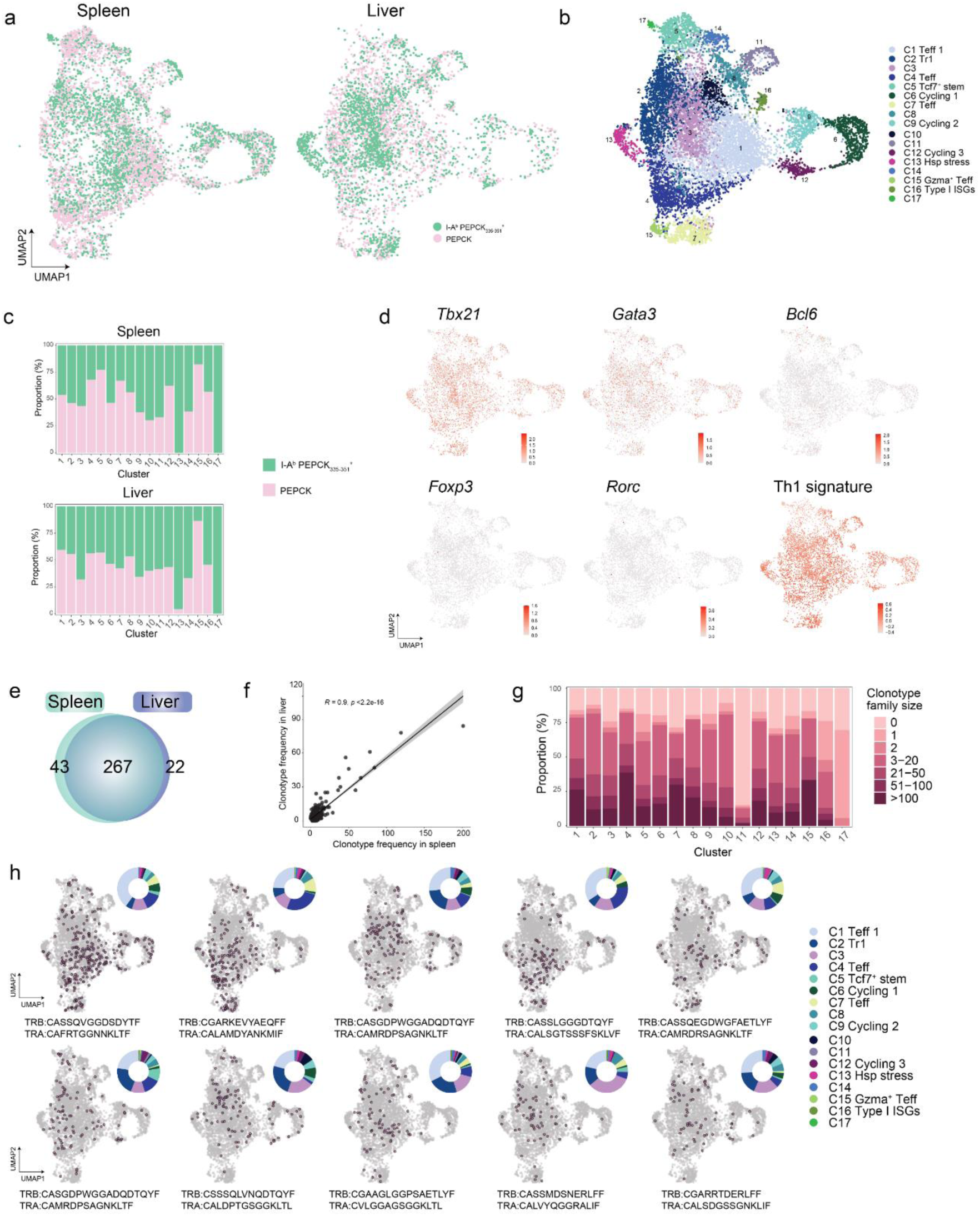
Leishmania-specific CD4^+^ T cell differentiation is independent of clonality or tissue localisation in experimental VL. **(a)** Droplet-based scRNAseq analysis of 10,696 PEPCK cells and I-A^b^ PEPCK_335-351_^+^ cells at day 14 p.i. in experimental VL. UMAP visualisation showing PEPCK and I-A^b^ PEPCK_335-351_^+^ cell type, subsetted by organ and **(b)** the clusters identified. **(c)** Bar graph shows the distribution of PEPCK cells and I-A^b^ PEPCK_335-351_^+^ cells from each organ across clusters. **(d)** UMAP visualisation showing subsetted I-A^b^ PEPCK_335-351_^+^ cells and the expression of master T helper cell transcription factors. **(e)** Venn diagram shows the distribution of 332 clonotypes identified with two or more family members in the I-A^b^ PEPCK_335-351_^+^ cells at day 14 p.i. between the spleen and liver. **(f)** Pearson’s correlation of the frequency of I-A^b^ PEPCK_335-351_^+^ cell clonotypes, with two or more family members, in the spleen and liver. **(g)** Bargraph showing the distribution of all I-A^b^ PEPCK_335-351_^+^ cell clonotypes in each cluster, grouped by family size. **(h)** The distribution of the top 10 most frequent I-A^b^ PEPCK_335-351_^+^ cell clonotypes across the UMAP visualisation for I-A^b^ PEPCK_335-351_^+^ cells only. Members of each clonotype are highlighted in pink. The doughnut plot shows the distribution of clonotype across clusters, colours correspond to clusters in (b).

We next subsetted I-A^b^ PEPCK_335-351_^+^ cells and examined the expression of lineage defining transcription factors. Consistent to the PEPCK cell phenotype during experimental VL, we observed high levels of *Tbx21* and Th1 cell gene signatures in I-A^b^ PEPCK_335-351_^+^ cells (Figure 3d). These data indicate that in experimental VL, the transcriptional landscape of TCR-restricted and polyclonal *Leishmania*-specific CD4^+^ T cells is conserved and closely aligns with Th1 cells.

### Tissue seeding and CD4^+^ T cell differentiation is largely independent of TCR diversity

Next, we investigated how CD4^+^ T cell clones are seeded between the spleen and liver to determine whether TCR diversity influences CD4^+^ T cell differentiation in experimental VL. In the I-A^b^ PEPCK_335-351_^+^ cells, we identified a TCR α– and β-chain for 4244 cells (79.36%) of 5348 cells total. Among these cells, we detected 583 clonotype families, defined by the sharing of identical TCR α– and β–chains, with 332 clonotypes that had a family size ≥ 2 cells. However, since the cells were pooled from multiple mice, it is important to note that clonotype diversity at the level of individual mice could not be assessed. To determine how CD4^+^ T cell clonotype influenced tissue distribution, we compared the localisation of the 332 clonotypes that had expanded. The majority of these clonotypes (80%) were found in PEPCK_335-351_^+^ cells isolated from both the spleen and liver, whereas only 43 and 22 clonotypes were unique to the spleen and liver, respectively (Figure 3e). We then compared the frequency of these 332 clonotypes in each tissue. Pearson correlation analysis identified a strong positive correlation indicating that in experimental VL, seeding of CD4^+^ T cell clonotypes in the spleen and liver occurs at a similar frequency (Figure 3f). To determine whether clonotype influences CD4^+^ T cell fate choices, we next examined how clonotypes were distributed between clusters. Although varying in frequency, clonotype family expansion was distributed across all clusters (Figure 3g), excluding Clusters 11 and 17 and no preference for a specific cluster was identified for the top 10 most expanded clonotypes (Figure 3h). Overall, these data indicate that TCR diversity had limited impact on CD4^+^ T cell tissue seeding or cluster differentiation in experimental VL.

### AmBisome treatment promotes hepatic tissue-resident memory-like CD4^+^ T cell development

The clustering analysis in Figure 2 identified Cluster 5 as a transcriptionally distinct, predominantly liver-resident subset that expanded following AmBisome treatment. These characteristics suggest a potential role for this CD4^+^ T cell subset in localised immune responses during experimental VL. To gain deeper insights into this subset, we performed a detailed analysis of its gene signatures and evaluated its functional relevance in the context of experimental VL.

Assessment of Cluster 5 marker genes (Figure 4a) led us to hypothesise that these cells were tissue-resident CD4^+^ T cells. We detected expression of *Zfp683*, *Cxcr6*, *Itga1*, *Cd69* and *Il7r* (Figure 4b), all of which have previously been linked to tissue-resident memory cells^27,28^. In addition, Cluster 5 was enriched for a CD8^+^ T cell Trm-gene signature^28^ (Extended Data Figure 6a), exhibiting upregulation of Trm-associated genes and downregulation of genes typically repressed in Trm cells. To confirm our transcriptional data corresponded with surface protein expression, we assessed the levels of CXCR6 and CD69 co-expression on PEPCK cells during experimental VL. At day 56 p.i., we identified a population of CXCR6^+^ CD69^+^ PEPCK cells in both the spleen and liver, with an average frequency almost two-fold greater in the liver. This difference became more pronounced following AmBisome treatment, resulting in an almost ten-fold increase in the liver, compared to the frequency observed in the spleen (Figure 4c & d). These data indicate that a subset of PEPCK cells acquires a Trm-like phenotype during experimental VL in both the spleen and liver, with AmBisome treatment promoting their accumulation in the liver.

**Figure 4.**
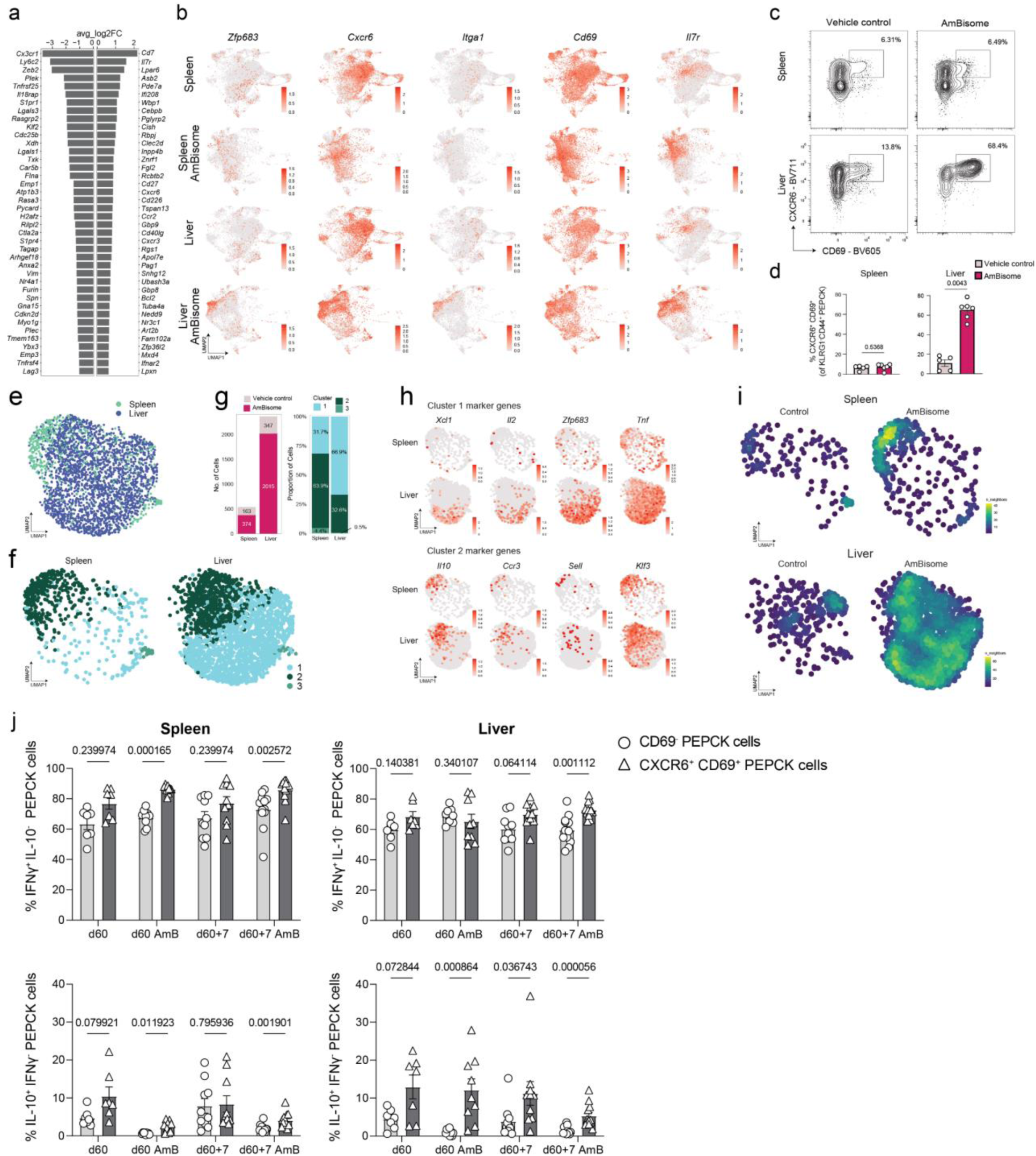
AmBisome treatment promotes the development of hepatic Trm-like CD4^+^ T cells. **(a)** UMAP visualisations of PEPCK cells subsetted by organ and treatment from d14-56 p.i. showing the expression of tissue-resident memory associated genes. **(b)** Waterfall plot showing the top 40 up- and down-regulated marker genes for Cluster 5, identified using the ‘FindAllMarkers’ function in Seurat. **(c)** Flow cytometry plots showing CXCR6 and CD69 expression on PEPCK cells (gated on KLRG1^-^ CD44^+^ PEPCK cells) from the spleen and liver at day 56 p.i. in experimental visceral leishmaniasis and **(d)** bar graphs showing the corresponding frequencies. Data is representative of two independent experiments (n=6 mice per experiment). Statistical testing was performed using two-tailed Mann-Whitney U test. Error bars represent mean ± SEM. **(e)** UMAP visualisation showing Cluster 5 after subsetting and data coloured by tissue of origin. **(f)** Three subsets of Trm-like PEPCK cells were identified based on unsupervised clustering. **(g)** Bargraphs show the total number of cells in Cluster 5 from the spleen and liver as well as the proportion of cells in each cluster for each tissue. **(h)** UMAP visualisations showing selected marker genes for Clusters 1 and 2. **(i)** Density plot showing the distribution of control and AmBisome treated samples for the spleen and liver. **(j)** Analysis of ex vivo IFNγ and IL-10 production by CD69^-^ and CXCR6^+^CD69^+^ PEPCK cells from the spleen and liver after PMA and ionomycin restimulation in vitro at day 60 and 7 days post-rechallenge (d60+7), in mice treated with vehicle control or AmBisome. Data are combined from two independent experiments (n=7 mice d60 control, n=9 mice d60 AmBisome, n=12 mice for d60+7). Error bars represent mean SEM. Statistical testing was performed using Multiple Mann-Whitney U tests..

We next compared the characteristics of Trm-like PEPCK cells in the spleen and liver to assess potential tissue-specific differences in their phenotype and function. Upon subclustering these cells, we observed distinct positional differences on the UMAP between spleen and liver PEPCK cells (Figure 4e). Clustering analysis revealed three clusters (Figure 4f), which were found in both tissues and across both control and AmBisome-treated mice, though their frequencies varied. Among splenic Trm-like PEPCK cells, the majority (64%) were in Cluster 2, while hepatic PEPCK cells were predominantly (67%) in Cluster 1. The marker genes for Cluster 1 included *Xcl1*, *Il2*, *Zfp683*, and *Tnf*, whereas Cluster 2 was enriched for *Il10*, *Ccr3*, *Sell*, and *Klf3* (Figure 4h). Additionally, we found that Ambisome treatment also affected the Trm-like cell phenotype (Figure 4i). Differential gene expression analysis revealed upregulation of exhaustion-associated genes in control Trm-like PEPCK cells, including *Lag3*, *Il21*, *Pdcd1*, and *Tigit*, while genes associated with enhanced function, such as *Il7r*, *Cxcr3*, *Cd226*, and *Ifnar2*, were upregulated in Trm-like PEPCK cells from AmBisome-treated mice (Extended Data Figure 6b). These data indicate that although Trm-like PEPCK cells develop in both the spleen and liver, there are distinct transcriptional differences between these cells in each tissue, with drug treatment further modulating their phenotype and gene expression profiles.

Next, we assessed the functional capacity of Trm-like PEPCK CD4⁺ T cells in the spleen and liver, and examined how this subset responded to antigen re-exposure. Following *in vitro* stimulation with PMA and ionomycin at day 56 post-infection, IFNγ production was comparable between CD69⁻ and CXCR6⁺CD69⁺ PEPCK cells in both tissues of vehicle control–treated mice (Figure 4j). In contrast, in AmBisome-treated mice, CXCR6⁺CD69⁺ PEPCK cells in the spleen produced significantly higher levels of IFNγ compared to their CD69⁻ counterparts prior to rechallenge. This enhanced IFNγ production by CXCR6⁺CD69⁺ cells persisted following rechallenge and was evident in both the spleen and liver. *In vitro* IL-10 production was significantly increased in CXCR6^+^CD69^+^ PEPCK cells from AmBisome-treated mice in both tissues, independent of co-inhibitory receptor or chemokine receptor expression (Extended Data Figure 6c). Together, these data indicate that CXCR6⁺CD69⁺ Trm-like CD4⁺ T cells exhibit enhanced functional potential compared to CD69⁻ cells, and are poised to produce IFNγ and IL-10 upon restimulation, especially in the liver, and this phenotype is promoted following drug treatment.

### *Bhlhe40* promotes effector CD4^+^ T cell function while suppressing CD4^+^ T cell stemness

The clustering analysis in Figure 2 suggested there were limited changes in the CD4^+^ T cell population structure dependent on tissue environment. However, to uncover gene expression patterns that were tissue-specific within clusters, differential gene expression analysis was performed between PEPCK cells from the spleen and liver at each time point in control-treated mice for the three largest clusters (Clusters 1 – 3). We identified 451, 391 and 282 differentially expressed genes in Cluster 1, 2 and 3, respectively (Extended Data Figure 8). These findings suggest that while the overall composition of CD4^+^ T cells is largely comparable between tissues, local microenvironmental factors may contribute to nuanced gene expression differences within specific CD4^+^ T cell subsets, potentially influencing functional specialisation and immune dynamics in VL.

One of the genes identified as consistently upregulated in PEPCK cells from the spleen compared to the liver in Clusters 1, 2 and 3 was *Bhlhe40*, a transcription factor known to regulate T cell differentiation and inflammatory responses^29^. Given its established role in regulating the production of IFN-γ and IL-10 in various infection models^30,31,32,33^ we focused on *Bhlhe40* to investigate its contribution to CD4^+^ T cell fate decisions in experimental VL. We first examined the role of *Bhlhe40* in regulating the production of IFNγ and IL-10 by CD4^+^ T cells *in vitro*. We employed CRISPR/Cas9 to target *Bhlhe40* in CD4^+^ T cells from triple reporter (*Il10^gfp^* x *Ifng^yfp^* x *Foxp3^rfp^*) C57BL/6 mice (Extended Data Figure 8a). Unstimulated CD4^+^ T cells were targeted with *Bhlhe40* sgRNA (*Bhlhe40*^KO^ CD4^+^ T cells), or a non-specific sgRNA as a control (*Bhlhe40*^WT^ CD4^+^ T cells), and then cultured under neutral (Th0), Th1 or Tr1 cell polarising conditions. Successful editing of *Bhlhe40* in *Bhlhe40*^KO^ CD4^+^ T cells was confirmed by RT-qPCR analysis. Targeting *Bhlhe40* did not effect the expression of *Maf* or *Prdm1*, genes associated with Tr1 cell differentiation^14,16^ (Extended Data Figure 8b), suggesting that the loss of *Bhlhe40* does not influence expression of these key regulatory transcription factors. However, when we assessed cytokine production, although we observed no significant change in IFNγ production, *Bhlhe40* deficiency caused a marked increase in IL-10 production in the Tr1-stimulation condition (Extended Data Figure 8c). Further, flow cytometry analysis revealed that the loss of *Bhlhe40* in CD4^+^ T cells resulted in an increased frequency of cells co-expressing both IFNγ and IL-10 (*IFNγ^+^IL-10^+^*), as well as a higher frequency of cells expressing IL-10 alone (*IFNγ^-^IL-10^+^*) under Tr1 cell conditions. In contrast, the frequency of IFNγ^+^IL-10^-^ CD4^+^ T cells was reduced in both the Th1 and Tr1 cell conditions.

To examine whether *Bhlhe40* played a role in determining tissue-specific outcomes of *L. donovani* infection, wild type (WT) and *Bhlhe40*-deficient (*Bhlhe40^-/-^*) mice were infected and parasite burdens and immune responses assessed in the spleen and liver at day 14 p.i.. In the absence of *Bhlhe40*, we observed a significant increase in parasite burdens in the liver and spleen, though the increase was more modest in the spleen (Figure 5a). Additionally, although spleen organ weight was significantly reduced in *Bhlhe40*^-/-^ mice (Extended Data Figure 8f), no differences in total cell numbers were detected in the spleen. In contrast, the liver of *Bhlhe40*^-/-^ mice showed a notable decrease in total cell numbers at day 14 p.i., suggesting that *Bhlhe40* may play an important role in regulating immune cell recruitment, effector functions and/or survival in the liver during infection.

**Figure 5.**
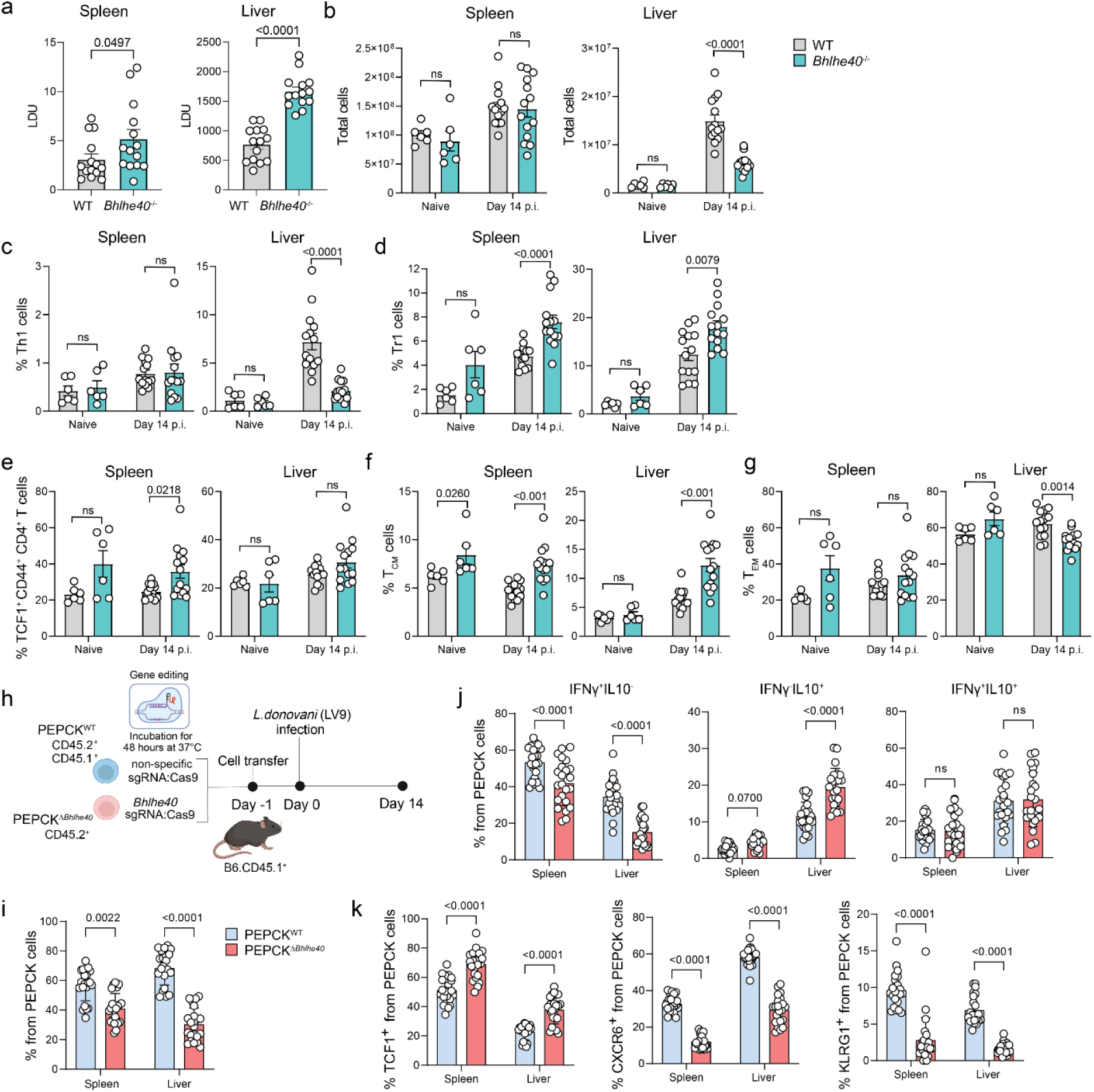
Bhlhe40 drives effector CD4^+^ T cell fate while restricting stemness in experimental VL. **(a)** Wildtype (WT) and Bhlhe40^-/-^ mice were infected with L. donovani and parasite burdens were measured in the spleen and liver at day 14 p.i.. LDU, Leishman-Donovan units. **(b)** Total number of cells in the livers of naïve mice and infected mice at day 14 p.i. **(c)** Frequency of Th1 cells (IFNƴ^+^Tbet^+^CD4^+^ T cells), **(d)** Tr1 cells (IFNƴ^+^IL-10^+^ CD4^+^ T cells), **(e)** TCF1^+^CD44^+^ CD4^+^ T cells, **(f)** central memory T cells (Tcm; CCR7^+^CD44^+^ CD4^+^ T cells), and (g) effector memory T cells (Tem; CCR7^-^CD44^+^ CD4^+^ T cells) in the spleen and liver of naïve and infected mice at day 14 p.i.. Statistical testing was performed using two-tailed Mann-Whitney U test. Error bars represent ± SEM. Data are pooled from two experiments, where n = 6 naïve WT and Bhlhe40^-/-^ mice, and n = 14 WT and Bhlhe40^-/-^ mice at day 14 p.i.. **(h)** CRISPR/cas9 editing of PEPCK cells targeting Bhlhe40 (PEPCK^ΔBhlhe40^), or a non-specific guide RNA (PEPCK^WT^), was performed and cells were subsequently co-transferred into B6.CD45.1^+^ mice, which were then infected with L. donovani. Flow cytometry assessment of PEPCK cells was performed at day 14 p.i.. **(i)** Graph showing the frequency of PEPCK^WT^ and PEPCK^ΔBhlhe40^ cells at day 14 p.i. from the spleen and liver. **(j)** Graphs showing the percentage of IFNƴ and IL-10 production by PEPCK^WT^ and PEPCK^ΔBhlhe40^ cells after in vitro stimulation with PMA and ionomycin at day 14 p.i. from the spleen and liver. **(k)** Graphs showing the expression of TCF1, CXCR6 and KLRG1 in PEPCK^WT^ and PEPCK^ΔBhlhe40^ cells at day 14 p.i. from the spleen and liver. **(i-k)** Statistical testing was performed using two-way ANOVA with Sidak’s multiple comparisons test. Error bars represent mean ± SEM. Data are pooled from four independent experiments, with n = 5/6 mice per group per experiment.

Next, we evaluated the impact of *Bhlhe40* deficiency on the development of Th1 (IFNγ^+^Tbet^+^) and Tr1 (IFNγ^+^IL-10^+^) cells in experimental VL. Compared to WT mice, *Bhlhe40*^-/-^ mice exhibited a significant reduction in the frequency of Th1 cells in the liver at day 14 p.i., while no such change was observed in the spleen (Figure 5c). In contrast, *Bhlhe40*^-/-^ mice displayed an increased frequency of Tr1 cells in both the spleen and liver at this time point (Figure 5d). As might be expected, these changes were reflected in significant alterations in the Th1:Tr1 cell ratios in both organs (Extended Data Figure 8g), suggesting that *Bhlhe40* plays a critical role in regulating the balance between Th1 and Tr1 cell responses during experimental VL. In *Bhlhe40*^-/-^ mice, we also observed enhanced TCF1^+^ CD44^+^ CD4^+^ T cell development in the spleen at day 14 p.i. (Figure 5e). *Bhlhe40* deficiency also promoted the development of central memory CD4^+^ T cells in both the spleen and liver at this time point (Figure 5f). Conversely, *Bhlhe40*^-/-^ mice exhibited a decrease in effector memory CD4^+^ T cell development in the liver (Figure 5g), supporting an important role for *Bhlhe40* in limiting the development of central memory and stem-like CD4^+^ T cells during experimental VL.

To establish cell intrinsic roles for *Bhlhe40* in CD4^+^ T cells during experimental VL, we performed CRISPR/Cas9 editing of PEPCK cells targeting *Bhlhe40* (PEPCK^Δ^*^Bhlhe40^*) (Figure 5h). Compared to control PEPCK (PEPCK^WT^) cells, the frequency of PEPCK^Δ^*^Bhlhe40^* cells was significantly reduced in both the spleen and liver, suggesting *Bhlhe40* may play a role in the expansion and/or survival of CD4^+^ T cells (Figure 5i). Flow cytometry analysis revealed a marked decrease in IFNγ⁺IL-10⁻ PEPCK cells in the absence of *Bhlhe40* in both organs, while IFNγ⁻IL-10⁺ PEPCK cells were significantly increased in the liver but remained unchanged in the spleen (Figure 5j). No differences were observed in the frequency of IFNγ⁺IL-10⁺ PEPCK cells. However, the loss of *Bhlhe40* resulted in a higher proportion of TCF1⁺ PEPCK cells in both the spleen and liver, accompanied by a reduction in CXCR6 and KLRG1 expression (Figure 5k). Together, these findings suggest that *Bhlhe40* regulates the balance between IFNγ and IL-10 production by CD4^+^ T cells, promotes effector cell differentiation and function, and suppresses either the development or maintenance of stem-like CD4^+^ T cells in experimental VL.

### *Tcf7*-expressing CD4^+^ T cells exhibit transcriptional characteristics associated with stemness in experimental VL

Stem-like CD4^+^ T cells represent an important memory cell population that reside in secondary lymphoid tissues, with a unique capacity to replenish effector T cells and sustain long-term immunity^34,35,36,37^. The clustering analysis in Figure 2 identified Cluster 7 as a spleen-specific CD4^+^ T cell subset characterised by *Tcf7* expression. Given the association of *Tcf7* with stem-like CD4^+^ T cells, we investigated whether this subset functions as a progenitor population capable of sustaining effector responses during chronic VL.

First, we examined the transcriptional and phenotypic characterisitics of Cluster 7 in more detail and observed expression of *Tcf7, Ccr7, Lef1, Cxcr5, Il7r,* and *Slamf6*, canonincal genes associated with stemness^34,38^ (Figure 6a). Several of these genes overlapped with the top marker genes that were differentially upregulated in this cluster (Figure 6b). In addition, when compared to a stem-like CD8^+^ T cell gene signature, we saw enriched expression of this signature in Cluster 7, and a corresponding downregulated effector CD8^+^ T cell gene signature (Extended Data Figure 9a). Together, these data suggest that Cluster 7 PEPCK cells exhibit a stem-like transcriptional profile.

**Figure 6.**
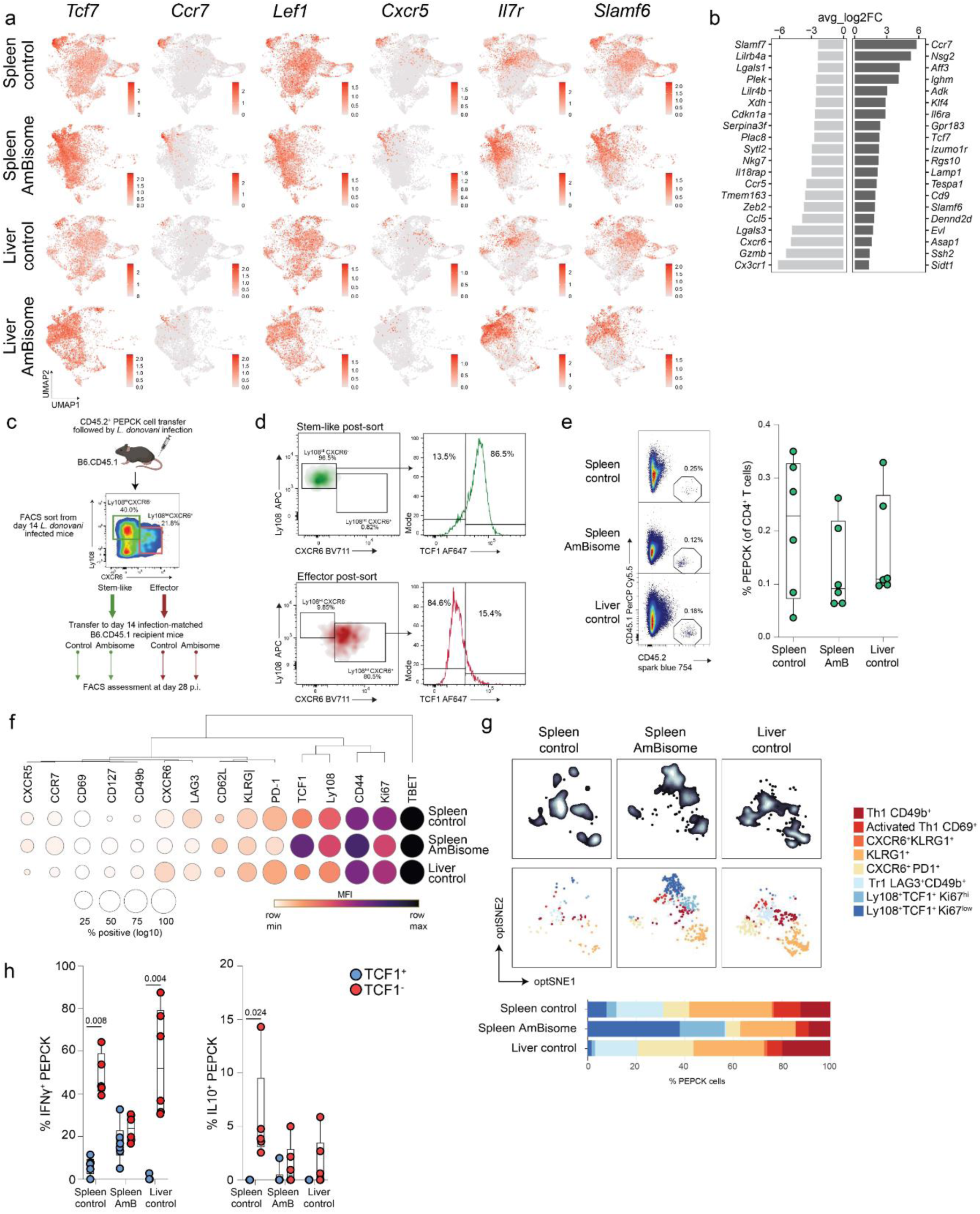
Tcf7-expressing CD4^+^ T cells exhibit stem-like properties in experimental VL. **(a)** UMAP visualisations of PEPCK cells subsetted by organ and treatment from d14-56 p.i. showing the expression of stemness-associated genes. **(b)** Waterfall plot showing the top 20 up- and down-regulated marker genes for Cluster 7, identified using the “FindAllMarkers()” function in Seurat. **(c)** Stem-like (Ly108^hi^Cxcr6^-^) and effector (Ly108^int^CXCR6^+^) PEPCK cells were isolated from the spleens of L. donovani infected mice at day 14 p.i. and adoptively transferred into infection matched mice that had been treated with vehicle control (control) or Ambisome (AmB) the day prior, followed by flow cytometry assessment at day 28 p.i.. **(d)** Flow cytometry plots showing the expression of TCF1 on isolated stem-like and effector PEPCK cells. **(e)** Representative flow cytometry plots and graph showing the frequency of PEPCK cells at day 28 p.i.. **(f)** The average expression (as MFI) and percentage (as positive events) is summarized in dot-plot format for PEPCK cells at day 28 p.i.. **(g)** optSNE visualisation showing PEPCK cell distribution and FlowSOM clustering (top) and the frequency of identified clusters for each sample (bottom). **(h)** Graphs showing the frequency of IFNƴ and IL-10 producing TCF1^+^ and TCF1^-^ PEPCK cells after in vitro stimulation with PMA and ionomycin. Statistical analysis was performed using multiple Mann-Whitney U tests. Data are from one experiment, with n = 5/6 mice per group.

Stem-like CD8^+^ T cells are a heterogeneous population^39^, so to examine whether similar diversity existed in stem-like PEPCK cells, we performed subclustering analysis. Three distinct clusters of stem-like PEPCK cells were identified (Extended Data Figure 9b), which exhibited a strong association with treatment type (Extended Data Figure 9c). At day 14 p.i., the majority of stem-like PEPCK cells originated from the spleen and were predominantly found in Cluster 2 (Extended Data Figure 9d). Beyond this time point, the number of splenic stem-like PEPCK cells declined significantly, except in AmBisome-treated mice where their frequencies remained stable. Consistent with this, flow cytometry analysis confirmed that the frequency of stem-like PEPCK cells in the spleen was maintained in AmBisome-treated mice, whereas they declined in vehicle-treated mice after day 14 p.i. (Extended Data Figure 9e). While these stem-like PEPCK cells persisted in the spleen following AmBisome treatment, their transcriptional profile shifted, with cells primarily transitioning to Cluster 1. Assessment of Cluster 1 marker genes identified those associated with self-renewal and longevity (*Il7r, S100a4, Klf6, Bcl2*)^40^ (Extended Data Figure 9f). In contrast, Cluster 2 marker genes were associated with activation and exhaustion (*Klrb1c, Il21, Lag3, Bcl6*), indicating a subset prone to functional decline. These data suggest that AmBisome treatment not only preserves splenic stem-like PEPCK cells but also supports their transition into a transcriptional state favoring long-term persistence rather than exhaustion. A feature of stem-like CD8^+^ T cells is their ability to preserve mitochondrial metabolism and promote long-term persistence via TGFβ- mediated mTORC1 inhibition^41^. Indeed, when mice with T cells lacking the ability receive TGFβ signals (B6.*Tgfbr2*^ΔT^) were infected with *L. donovani*, there were significant reductions in TCF1 expression and frequencies of stem-like CD4^+^ T cells at day 14 p.i., compared with littermate control mice (Extended Data Figure 10a). Furthermore, there was reduced expression of activation markers such as CXCR6, KLRG1 and IFNγ by B6.*Tgfbr2*^ΔT^ CD4^+^ T cells in the liver (Extended Data Figure 10b), suggesting similar metabolic requirements for the development of stem-like CD4^+^ T cells.

Given the similarities between stem-like T cells and central memory T cells (Tcm), we examined whether there was overlap between stem-like PEPCK cell clusters and Tcm cells. Classical Tcm cells express *Sell* and *Ccr7*, enabling lymphoid circulation, but most stem-like PEPCK cells lacked *Sell* expression (Extended Data Figure 9g), suggesting they do not fully align with a Tcm identity. Although Cluster 3 showed the closest resemblance to Tcm due to its expression of *Ccr7* and Sell, this cluster constituted only ∼2% of the total stem-like population, indicating that these cells are rare, and their low frequency makes definitive classification as conventional Tcm challenging.

### Splenic stem-like CD4^+^ T cells help sustain effector responses during chronic infection in experimental VL

To further investigate the role of *Tcf7* in CD4^+^ T cells, we used CRISPR/Cas9-mediated gene editing to disrupt *Tcf7* in PEPCK cells. Wild-type (PEPCK^WT^) and *Tcf7*-edited (PEPCK^Δ^*^Tcf7^*) PEPCK cells were co-transferred into recipient mice, which were subsequently infected with *L. donovani* (Extended Data Figure 9h). Fourteen days post-infection, we assessed the impact of *Tcf7* deletion on PEPCK cell persistence and phenotype using flow cytometry. Surprisingly, we observed no significant differences between PEPCK^WT^ and PEPCK^Δ^*^Tcf7^* cells in terms of frequency or survival (Extended Data Figure 8i). While most surface marker expression remained unchanged between PEPCK^WT^ and PEPCK^Δ^*^Tcf7^* cells, we did observe slightly increased CXCR6 expression, and reduced KLRG1 expression in PEPCK^Δ^*^Tcf7^* cells compared to the PEPCK^WT^ cells (Extended Data Figure 8j). However, by knocking out *Tcf7* prior to infection, we may have allowed *Tcf7*-independent pathways to emerge to drive CD4^+^ T cell expansion and effector responses.

Therefore, to directly assess the self-renewal potential and differentiation capacity of stem-like CD4^+^ T cells, we employed an adoptive transfer model using stem-like and effector PEPCK cells. For this, we transferred PEPCK cells (CD45.2^+)^ into B6.CD45.1^+^ mice and subsequently infected them with *L. donovani*. On day 14 p.i., when splenic stem-like PEPCK cells had established, we isolated stem-like (Ly108^hi^Cxcr6^-^), where we used Ly108 as a surrogate for TCF1 expression^37^, and effector (Ly108^int^Cxcr6^+^) PEPCK cells from the spleens of these mice (Figure 6c). The expression of TCF1 by stem-like PEPCK cells was confirmed by flow cytometry (Figure 6d) prior to adoptively transferring both stem-like and effector PEPCK cells into separate groups of infection matched B6.CD45.1 recipient mice that had either been treated with vehicle control or AmBisome on the day prior. After 14 days, we then performed flow cytometry assessment of PEPCK cells from these recipient mice. PEPCK cells were identified in the spleens and livers of mice that had received stem-like PEPCK cells, except in the livers of AmBisome-treated mice, where only few PEPCK cell numbers were observed (Figure 6e). In contrast, no PEPCK cells were detected in the spleen or liver of mice that had received the effector PEPCK cells. These findings suggest that TCF1 expression is essential for the self-renewal and maintenance of these cells, which in turn, differentiate into effector CD4^+^ T cells.

Surface marker analysis confirmed that PEPCK cells from all three groups where they could be readily detected, exhibited a Th1 cell phenotype, marked by Tbet expression, and were actively proliferating, as indicated by Ki67 expression (Figure 6f). In both the spleen and liver, PEPCK cells from vehicle control-treated mice displayed higher expression of Lag3 and PD-1 compared to those from AmBisome-treated mice. Notably, splenic PEPCK cells from AmBisome-treated mice retained a stem-like phenotype, characterised by TCF1 expression. To further define these populations, unsupervised clustering of PEPCK cell FACS data identified eight distinct clusters (Figure 6g). These included five effector-like clusters (Cd49b^+^, CD69^+^, CXCR6^+^KLRG1^+^, KLRG1^+^ and CXCR6^+^PD1^+^), a regulatory cluster (Tr1; LAG3^+^CD49b^+^) and two stem-like clusters (Ly108^+^TCF1^+^Ki67^hi^, Ly108^+^TCF1^+^Ki67^low^), distinguished by their Ki67 expression levels. While all clusters were present across treatment groups, their frequencies differed (Figure 6g). Notably, AmBisome-treated mice exhibited an increased proportion of splenic stem-like PEPCK cells compared to vehicle control treated mice, with a significant enrichment of the Ly108^+^TCF1^+^Ki67^low^ subset, indicative of a more quiescent phenotype. Consistent with this, functional analysis of PEPCK cells following *in vitro* stimulation with PMA and ionomycin revealed that TCF1^+^ PEPCK cells produced little to no IFNγ or IL-10, whereas TCF1^-^ PEPCK cells readily produced these cytokines (Figure 6h). These findings suggest that stem-like PEPCK cells in the spleen remain in a functionally quiescent state, with AmBisome treatment promoting their maintenance as a long-term reservoir that may sustain immune responses in experimental VL.

Finally, to test whether stem-like PEPCK cells can give rise to anti-parasitic CD4^+^ T cells, we isolated stem-like (Ly108^hi^Cxcr6^-^) and effector (Ly108^int^Cxcr6^+^) PEPCK cells from the spleens, as above, and transferred them into B6.*Rag1*^-/-^ mice the day prior to *L. donovani* infection. We found limited impact of either population on the control parasite growth in the spleen, as expected given the relatively low parasite burdens at this early timepoint, but a similar ability of both populations to control parasite growth in the liver (Extended Data Figure 11). Thus, stem-like CD4^+^ T cells reside in the spleen in a functionally quiescent state, but retain their ability to differentiate into effective anti-parasitic CD4^+^ T cells.

## Discussion

Effective CD4^+^ T cell responses are critical for immunity to VL, yet the factors shaping their differentiation and function across distinct tissue environments remain poorly understood. Here, we explored how CD4^+^ T cells adapt to the contrasting immune landscapes of the spleen and liver during experimental VL, uncovering key differences in their composition, transcriptional profiles and functional states. By integrating single-cell transcriptomics with complementary analyses, we captured the heterogeneity of these responses and identified key transcriptional programs associated with effector function, regulatory potential and stem-like properties. Furthermore, we examined how anti-parasitic treatment with AmBisome influences CD4^+^ T cell dynamics, shedding light on mechanisms that likely contribute to long-term immunity in VL. Together, these findings offer a detailed framework for understanding tissue-specific immunity in VL and provide insights relevant to both therapeutic interventions and vaccine development.

CD4^+^ T cells are essential for resolving *L. donovani* infection, with the balance between Th1 and Tr1 cells often considered a key determinant of disease outcome^42^. However, our study provides the first in-depth, high-resolution analysis of CD4^+^ T cell responses in experimental VL, revealing that CD4^+^ T cell heterogeneity extends far beyond this conventional Th1-Tr1 cell framework. Rather than being a simple binary developmental pathway, we uncovered a diverse spectrum of transcriptional states that shape immunity and disease progression in a more nuanced and tissue-specific manner. We identified multiple Th1 cell subtypes, including effector, memory-like, regulatory and stem-like subsets, underscoring a broader spectrum of Th1 cell functional states than previously appreciated. While the classification of CD4^+^ T cells into distinct subsets such as Th1, Th2, Th17, Tfh and Treg cells is well established^10^, it is increasingly recognised that heterogeneity exisits within these broad categories. For example, studies on Th17 and Tfh cells have highlighted the existence of distinct functional subpopulations with different transcriptional programs and capacities^43,44^. However, the diversity within the Th1 cell lineage has remained less well explored. CD4^+^ T cells exhibit phenotypic plasticity, often co-expressing transcriptional programs associated with multiple cell subsets^45,46^, and our findings suggest that such flexibility extends to Th1 cells in the context of VL. This highlights the need to consider how different Th1 cell subsets, and their balance, contribute to immune responses in chronic infection settings.

The similarity between polyclonal *Leishmania*-specific CD4^+^ T cells and monoclonal PEPCK cells in our study suggests that TCR diversity is not a major driver of tissue localisation or differentiation patterns in experimental VL. This observation aligns with findings in other chronic infections, such as lymphocytic choriomeningitis virus (LCMV)^47,48^, where CD4^+^ T cells exhibit functional plasticity irrespective of their TCR specificity. These insights underscore the significance of tissue-specific microenvironmental factors, such as cytokine milieu, stromal interactions, and metabolic conditions in guiding CD4^+^ T cell differentiation and localisation. Consequently, therapeutic strategies that modulate these extrinsic factors may be more effective in shaping protective immunity in VL.

Our analysis revealed striking transcriptional similarities between CD4^+^ T cells from the spleen and liver, challenging our initial hypothesis that intrinsic differences in these cells drive the disparity in parasite control between these organs in experimental VL. Instead, our findings suggest that extrinsic factors within the tissue microenvironment likely play a more dominant role. The spleen undergoes profound structural changes during chronic infection, including splenomegaly and loss of lymphoid tissue organisation^17,18,19^. In contrast, the liver mounts a granulomatous response that effectively contains infection. This breakdown in splenic architecture, driven in part by excessive TNF signaling^49^, likely contributes to the failure of parasite control in the spleen despite the presence of transcriptionally similar CD4^+^ T cell populations in both organs. Understanding the localisation of CD4^+^ T cell subsets within the spleen during infection could provide key insights into how their interactions with other immune cells and stromal networks become disrupted, ultimately impairing their ability to effectively clear parasites.

Although broad transcriptional differences between the spleen and liver were limited, we identified shifts in subset frequencies, including the presence of a novel stem-like CD4^+^ T cell population in the spleen. Stem-like T cells support sustained immune responses and retain the capacity to differentiate into diverse effector subsets^34,36^. However, *Bhlhe40*, identified as upregulated in the spleen, emerged as a key suppressor of this phenotype. *Bhlhe40* deficiency skewed CD4^+^ T cell responses, reducing Th1 cell differentiation while promoting Tr1 cells, as has previously been reported in *Toxoplasma gondii* and *Mycobacterium tuberculosis* infection^30,33^. This also correlated with impaired parasite control in the liver and spleen in experimental VL. Additionally, *Bhlhe40* restricted TCF1 expression, thereby influencing stem-like CD4^+^ T cell development. This raises the possibility that *Bhlhe40* regulates the persistence and function of TCF1⁺ PEPCK cells, with broader implications for T cell fate decisions in chronic infection. These findings highlight the complexity of CD4^+^ T cell differentiation in VL, suggesting that tissue-specific cues shape not only effector responses but also the balance between renewal and exhaustion.

The most profound transcriptional shift in CD4^+^ T cells occurred following AmBisome treatment, an observation with important implications for VL immunity. In human VL, patients who recover often exhibit long-term resistance to reinfection, highlighting the need to understand how the immune landscape is reshaped post-treatment. In our model, AmBisome treatment preserved the stem-like CD4^+^ T cell population while also promoting the development of Trm-like CD4^+^ T cells. Trm cells provide rapid localised immune responses upon pathogen re-exposure^50^, and their presence likely contributes to effective immunity against *L. donovani* infection. Defining the roles of these populations in VL immunity will be critical for guiding future therapeutic and vaccine strategies. Understanding how treatments like AmBisome modulate the immune landscape can inform the development of interventions that not only clear the infection but also establish robust, long-lasting immunity.

Together, our findings provide a comprehensive view of the complexity of CD4^+^ T cell responses in experimental VL, revealing a previously unappreciated heterogeneity in Th1 cell subsets and highlighting the critical influence of the tissue microenvironment, infection stage, and drug treatment on their differentiation and function. The observation that CD4^+^ T cells from the spleen and liver share highly similar transcriptional profiles challenges traditional assumptions about organ-specific immune programming and suggests that extrinsic factors, rather than intrinsic differences in T cells, drive the disparate control of *Leishmania donovani* between these tissues. The identification of stem-like PEPCK cells and their persistence following drug treatment raises intriguing questions about their role in long-term immunity and whether they can be harnessed for therapeutic benefit. Furthermore, the promotion of Trm-like cell populations post-treatment underscores the potential for tissue-resident memory responses to contribute to durable immunity in VL. The involvement of transcriptional regulators such as Bhlhe40 in shaping these immune dynamics presents an exciting avenue for future research, particularly in understanding how interventions might be tailored to optimise protective immunity while preventing immune exhaustion. More broadly, these findings have implications beyond VL, contributing to our understanding of how CD4^+^ T cell heterogeneity and plasticity shape immune responses in chronic infections. By dissecting the mechanisms that govern CD4^+^ T cell fate decisions, we can refine immunotherapeutic strategies not only for leishmaniasis but also for other persistent infections, autoimmune conditions, and vaccine development.

## Materials and methods

### Mice

B6.CD45.1^+^ (B6.SJL-Ptprc^a^ Pepc^b^/BoyJ) mice were purchased from Animal Resources Centre/ Ozgene (Canning Vale), Bhlhe40^-/-^ mice (B6.129S1(Cg)-*Bhlhe40^tm1.1Rhli^*/MpmJ) were purchased from The Jackson Laboratory, and C57BL/6J.PEPCK and B6.Rag1^-/-^ mice were bred in house. C57BL/6-Il10tm1Flv/J (*Il10^gfp^*; JAX:008379)^51^, B6.129S4-Ifngtm3.1Lky/J (*Ifng^yfp,^* RRID: IMSR_JAX: 017581)^52^ and C57BL/6-*Foxp3^tm1fl^v*/J (*Foxp3^rfp^*; JAX:008374) mice were crossed to produce a triple reporter strain. C57BL/6-129-Tgfbr2^tm1karl^/J (*Tgfbr2^fl/fl^*; JAX:012603)^53^ were crossed with C57BL/6-Cg-Tg(Lck-icre)3779Nik/J (dLck^cre^; JAX:012837)^54^ to produce mice with T cells lacking TGFβRII (B6.*Tgfbr2*^ΔT^). The Cre-negative littermates were used as controls. All mice were female, aged 8-12 weeks, and maintained under specific pathogen-free conditions within the animal facility at QIMR Berghofer. Mice were housed in exhaust-ventilated cages (Opti-mice) with ≤6 mice per cage, and room temperature was maintained between 19 °C and 22 °C, with humidity between 55% and 65% and with 12-h/12-h dark/light cycle with a 15-min sunrise and sunset. All animal procedures and protocols were in accordance with the “Australian Code of Practice for the Care and Use of Animals for Scientific Purposes” (Australian National Health and Medical Research Council [NHMRC]) and approved by the QIMR Berghofer Animal Ethics Committee (Brisbane, Queensland, Australia; approval number A1707-615M/A2305-606 or P2304/P3887).

### PEPCK adoptive transfer

Spleens were collected from C57BL/6J.PEPCK mice and homogenised through a 100 µm cell strainer to create a single cell suspension. Red blood cells were lysed using Red Blood cell Lysing Buffer Hybri-Max (Sigma-Aldrich) and samples were washed with 1% FCS/PBS. Cells were enriched for CD4^+^ T cells using EasySep™ Mouse CD4^+^ T Cell Isolation kits (Stemcell Technologies), according to the manufacturer’s protocol. Naïve PEPCK cells (CD62L^hi^ CD44^lo^) were isolated by flow cytometry and then transferred to each mouse (1 x 10^4^ cells in 200 µl volume per mouse) via lateral tail vein intravenous (i.v.) injection.

### Leishmania donovani infections in mice

*Leishmania donovani* (LV9; MHOM/ET/67/HU3) parasites were maintained by passage in B6.Rag1^-/-^ mice and amastigotes were isolated from the spleens of chronically infected mice, as previously described^26^. Mice were infected with 2 x 10^7^ LV9 amastigotes i.v. (in 200 µl volume) via the lateral tail vein.

### Quantifying *Leishmania* parasite burdens

Spleen and liver impression smears were used to determine parasite burdens and were expressed as Leishman Donovan Units (LDU; number of amastigotes per 1000 host nuclei multiplied by the organ weight (in grams)).

### AmBisome drug treatment

Liposomal amphotericin B (AmBisome^®^; Gilead Sciences) was prepared according to the manufacturer’s instructions. Mice were treated with a single dose of AmBisome (10 μg/g body weight) or vehicle control (3.875% glucose (vol/vol)) at day 14 post-infection, administered i.v. (200 µl volume per mouse) via the lateral tail vein.

### Preparation of spleen and liver single-cell suspensions

Spleens were collected and homogenised through a 100-μm cell strainer to create a single cell suspension. Red blood cells were lysed using Red Blood cell Lysing Buffer Hybri-Max (Sigma-Aldrich) and samples were washed with 1% FCS/PBS. Cells were resuspended in 1% FCS.PBS and counted using a Countess Cell Counting Chamber Slides on the Countess II FL (Invitrogen), per the manufacturer’s protocol. Cells were enriched for CD4^+^ T cells using EasySep™ Mouse CD4^+^ T Cell Isolation kits (Stemcell Technologies) or magnetic-activated cell sorting (MACS) using anti-mouse CD4 Microbeads (Miltenyi Biotec), according to the manufacturer’s protocol.

Livers were first perfused via the hepatic portal vein with 1x PBS and then homogenized through a 100-μm cell strainer to create a single cell suspension. Hepatocytes were separated from leukocytes using a 33% (vol/vol) Percoll Density Gradient (GE Healthcare). After removal of supernatant containing unwanted cells and debris, the liver leukocyte pellet was processed as described above for the splenic tissue.

### Flow cytometry

LIVE/DEAD™ Fixable Blue Dead Cell Stain Kit, for UV excitation (ThermoFisher Scientific) was used as the initial step to exclude dead cells. Cells were then incubated with 0.2µl/sample TruStain FcX™ PLUS (anti-mouse CD16/32) antibody (Biolegend) for blocking of Fc receptors, CCR7-PeCy7 (Biolegend) was stained individually for 15 minutes at 37°C in the dark followed by surface staining with a combination of the following antibodies; CD44 AF-700 or FITC (Biolegend), CD90.2 PerCP-Cy5.5 (Biolegend), CD127 BV421 (Biolegend), CD69 BV605 (Biolegend), CD62L BV650 or PE (Biolegend), CXCR6 BV711 (Biolegend), CD45.2 Sparkblue-754 or PerCP-Cy5.5 (Biolegend), CD45.1 PerCP-Cy5.5 (Biolegend), KLRG1 PEDazzle-594 (Biolegend), PD1 APCFire-750 (Biolegend), Ly108 APC (Biolegend), CD4 BUV395 (BD Biosciences) or AF647 (Biolegend), CXCR5 BV785 and TCRβ BUV737 (BD Biosciences), for 60 minutes at 37°C in the dark.

For intracellular cytokine staining, cells were first stained with a mix of LAG3 BV785, CCR2 BV421, CCR5 PerCP-Cy5.5 and CD49b PeCy7 for 30 minutes at 37°C in the dark and then stimulated with a combination of 50ng/ml Phorbol-12-myristate-13-acetate (PMA; Sigma Aldrich) and 1uM Ionomycin (Sigma Aldrich) for 3 hours at 37°C in the presence of 1x Monensin (1000x Biolegend) or 1x Brefeldin A (1000x Biolegend) followed by surface staining as above. Cells were then fixed and permeabilised with the eBioscience™ Intracellular Fixation and Permeabilisation Buffer Set (Thermofisher), according to the manufacturers’ instructions, followed by staining for IL10 PE-Dazzle594 (Biolegend), IFNƴ BV650 (BD), Foxp3 Ef450 (eBioscience), TCF1 PE (Cell Signaling Technolgies), Tbet PE-Cy7 (Biolegend), Ki67 APC-fire 750 (Biolegend), CTLA4 BV650 (Biolegend) for 45 minutes at 37°C in the dark.

Data were acquired using a Cytek® Aurora spectral analyser (Cytek Biosciences). Data were gated using FlowJo software (TreeStar) and analyzed with GraphPad Prism v9. For clustering analysis the OMIQ™ (https://www.omiq.ai/) platform was used to batch correct data with gaussNorm, followed by tSNE clustering and FlowSOM analysis to identify unique clusters. Mean fluorescent intensity heat maps and expression dot plots were made using MORPHEUS (https://software.broadinstitute.org/Morpheus) (Broad Institute).

### Tetramer staining

*Leishmania*-specific CD4^+^ T cells were identified by staining with a MHC-class II I-A^b^ PEPCK_335-351_ APC-conjugated tetramer. To minimise unspecific binding of cells, cells were also stained with a control human CLIP (I-A^b^ CLIP_87-101_) phycoerythrin (PE)– conjugated tetramer. The tetramers were produced by the NIH Tetramer Core Facility and cells were stained for 30 minutes at 37°C.

### Single-cell RNA capture and sequencing

Single cell suspensions from mouse spleen and liver were prepared as described above. Cells were then incubated with TruStain FcX™ PLUS (anti-mouse CD16/32) antibody (BioLegend) for blocking of Fc receptors followed by surface staining with a combination of surface antibodies (as listed in Supplementary Table 1) for 30 minutes on ice in the dark. In addition, samples were also incubated with TotalSeq™-C anti-mouse Hashtag antibodies (Biolegend; Supplementary Table 1) on ice for 30 minutes in the dark and washed with 1% BSA.PBS. Cells were then resuspended in 1% BSA.PBS containing SYTOX™ Blue Dead Cell Stain (ThermoFisher Scientific) at a dilution of 1:1000. PEPCK cells (CD4^+^CD90.2^+^CD45.2^+^CD45.1^-^) were then sorted using a FACS ARIA III (BD) into a 1% BSA.PBS buffer. After sorting, PEPCK cells were loaded onto a Chromium Single Cell Chip (10x Genomics) for generation of gel bead-in emulsions (GEMs), according to manufacturers’ instructions, with a target capture rate of ∼4000 single cells per sample. Generation of cDNA and single-cell RNA sequencing libraries for Illumina sequencing were generated using the Chromium Single Cell 5’ v2 kit (10x Genomics). All samples from a single time point were processed together in one well of the Chromium chip and the resulting libraries were prepared in parallel. All libraries were pooled and sequenced on five Illumina flow cells with a 10-base index read, a 26-base read 1 containing cell-identifying barcodes and unique molecular identifiers (UMIs), and a 90-base read 2 containing transcript sequences on an Illumina NextSeq550.

### Single-cell RNA sequencing data processing and analysis

Illumina FASTQ files were run through the ‘cellranger count’ pipeline from Cell Ranger v6.0.1 (10x Genomics) with 10x Genomics mouse genome mm10-2020-A as a reference. *Seurat* v4^55^ was used for downstream analysis. Only genes expressed in three or more cells were considered. Cells were filtered to remove those expressing fewer than 200 genes and more than 5000 genes, and those with more than 10% mitochondrial content. Cells identified as doublets were removed by demultiplexing based on HTO using “HTODemux()” function from *Seurat* package. To minimise TCR-driven variability in clustering, T cell receptor alpha and T cell receptor beta genes were removed prior to downstream analysis. Additionally, a cluster of contaminating hepatocytes was excluded from the dataset.

To perform principal component analysis (PCA), we normalised the data using “SCTransform()” function from *Seurat* package with default settings, then used this as input for the “RunPCA()” function from *Seurat* package. The computed principle components (PCs) were used as input to generate uniform manifold approximation and projection (UMAP) using “RunUMAP()” from *Seurat* package. Unsupervised clustering was performed using “FindNeighbours()” function, followed by “FindClusters()” function from *Seurat* package. Gene signature scoring was performed by computing signature scores for each cell in the dataset using “AddModuleScore()” function from *Seurat* package. Differential gene expression analysis was performed using “FindMarkers()” function from *Seurat* package. Genes with adjusted *P*-values <0.05 were considered as differentially expressed genes.

### Nucleofection

CRISPR Cas9 gene editing was performed on unstimulated mouse CD4^+^ T cells. Mouse splenic CD4^+^ T cells were isolated using an EasySep Mouse CD4^+^ T cell Enrichment Kit (STEMCELL Technologies), according to the manufacturers’ instructions. A 3:1 ratio of Cas9-RNP complex was prepared by mixing 2μl sgRNA (120pmol) (Synthego) and 2μl Cas9 (40pmol) (QB3 Macrolab) followed by incubation at 37°C for 15 minutes. Multi-guides used for each gene were; *Bhlhe40*: A*A*A*CUUACAAACUGCCGCAC, C*G*A*GUGCAUUGCCCAGCUGA, U*C*U*GGGCAGAAGAACUGGGU; *Tcf7*: U*A*A*GAAUCGCGACAGCGCUG, G*C*A*GCUCGUCGGGCGCGCCG, A*U*G*CCGCAGCUGGACUCGGG. Using Amaxa™ P4 primary cell 96-well Nucleofection™ kit (Lonza), 1-5×10^6^ CD4^+^ T cells in 20μL P4 buffer was added to the 2μL of RNP mixture, and the total 24μL mixture was transferred to a nucleofection well strip. Nucleofection was performed using the Amaxa™ Nucleofector 96-well shuttle system (Lonza), selecting program DS137. Afterwards, 100μL of 37°C T cell culture media (cRPMI, see *Supplementary 8.1. General reagents*) was added to cell samples and transferred to a 96 U-bottom well plate, followed by addition of 125μL cRPMI. 1μL of 2500ng/mL recombinant mouse IL-7 and 2.5μL of 200μg/mL IL-2 was then added to each well and incubated for 48 hours at 37°C in 5% (v/v) CO2. For *in vivo* functional studies, a 1×10^6^ cells/200μL dPBS mix containing equal amounts, confirmed by flow cytometry, of CRISPR/Cas9 gene edited and negative control naïve CD4^+^ T cells were intravenously injected into B6.CD45.1^+^ recipient mice via a lateral tail vein.

### *In vitro* CD4^+^ T cell polarisation

For *in vitro* polarisation, a 96 flat-well plate was coated with 100μL of 5μg/mL anti-CD3ε mAb (Biolegend). CRISPR/Cas9 gene edited and negative control CD4^+^ T cells (2.5×10^5^) supplemented with 5μg/mL anti-CD28 mAb (Biolegend) and 2.5ng/ml recombinant mouse IL-2 were transferred to the anti-CD3ε-coated flat-well plate, and incubated for 72 hours at 37°C in 5% (v/v) CO2 in the presence of polarising antibodies and cytokines in a final volume of in 250μL of cRPMI. For Th1 polarisation: 20ng/mL IL-12 (Biolegend), and 20µg/mL αIL-4 mAb (Biolegend); Tr1 polarisation: 20ng/mL IL-12, 200ng/mL IL-27 (Prepotech), and 20µg/mL αIL-4 mAb. Cytokine levels from cell culture supernatants were measured by flow cytometry using the Cytometric Bead Array mouse Th1/Th2/Th17 Cytokine kit (BD Biosciences) as per manufacturer’s instructions on an LSRFortessa™ (BD Biosciences), and analysed using FCAP Array™ Software v3.0 (BD Biosciences).

### Real-time quantitative PCR

RNA was extracted from CD4^+^ T cells with the RNeasy Mini Kit (Qiagen) following manufacturer’s instructions, RNA was reverse transcribed to cDNA using the ZymoScript RT PreMix Kit (Zymoe) and real-time qPCR was performed with the Gotaq probe Kit (Promega) for mouse *Bhlhe40, Maf* and *Prdm1* on a Bio-Rad CFX 384 real-time PCR system (Bio-Rad) using the TaqMan Gene Expression Assay for *Bhlhe40* (Mm00478592_m1; Applied Biosystems), *Maf* (Mm02581355_s1; Applied Biosystems) and *Prdm1* (Mm00476128_m1; Applied Biosystems). Relative expression was normalised relative to mouse *Hprt* (Mm03024075_m1; Applied Biosystems) housekeeping gene.

### Cell sorting and adoptive transfer of stem-like and effector PEPCK cells

Spleen mononuclear cells from mice at day 14 of infection with *L. donovani* were stained with CD45.1 FITC (Biolegend), Ly108 APC (Biolegend), CD45.2 PE (Biolegend), CD4 BV605 (Biolegend) and CXCR6 BV711 (Biolegend) for 30 minutes at 37°C in the dark followed by SYTOX™ Blue Dead Cell Stain (ThermoFisher Scientific). Cell sorting was performed on a BD FACSAria™ III (BD Biosciences) instrument, with purity confirmed at 80-95%. Purified cells were then injected via tail intravenous injection at 2.5×10^4^ cells for Ly108^-^CXCR6^+^ PEPCK cells or 9×10^4^ for Ly108^+^CXCR6^-^ PEPCK cells into infected congenic (B6.CD45.1) mice. For the B6.*Rag1*^-/-^ adoptive transfer experiments, 3 x 10^4^ Ly108^-^CXCR6^+^ PEPCK cells or Ly108^+^CXCR6^-^ PEPCK cells were transferred into mice one day prior to infection.

### Statistical analysis

Statistical analyses were performed using Prism 9 (version 9.4.0; GraphPad Software). Statistical testing was performed using either the Mann-Whitney *U*-test or two-ANOVA with Sidak’s multiple comparisons test, unless stated otherwise.

## Data and materials availability

The materials, data and any associated protocols that support the findings of this study are available from the corresponding author upon request. The raw sequencing data and processed Cell Ranger outputs used in this study will be deposited in an online repository on publication.

## Author contributions

JAE, FDLR, BC, JN, PTB, LB, TCMF and YW generated data, supervised by JAE, FDLR, AH or CRE. JAE, FDLR, BC, SA and HJL analysed data. JAE, AH and CRE conceptualised the study. KC, KHG provided reagents. JAE and CRE led manuscript writing. All authors contributed to and approved the manuscript.

## Acknowledgements

We thank the QIMR Berghofer Flow Cytometry facility, Animal House facility, DNA Sequencing facility, and Genome Informatics Group for their support. We thank the NIH tetramer facility (Atlanta, GA, United States) for production of the I-A^b^-PEPCK_335–351_ tetramer used to detect *L. donovani* PEPCK-specific CD4^+^ T cells and I-A^b^-CLIP_87-101_ tetramer in these studies. This work was supported by the National Health and Medical Research Council of Australia; Program Grant (1132975) to C.R.E, Senior Research Fellowships to C.R.E (1154265), Australian Infectious Diseases Research Centre University of Queensland-QIMR Berghofer Collaborative Research Scheme to C. R. E. and J. A.E., and QIMR Berghofer Seed Funding Scheme to J.A.E.

## Extended Data Figures

**Extended Data Figure 1 (related to Figure 1).**
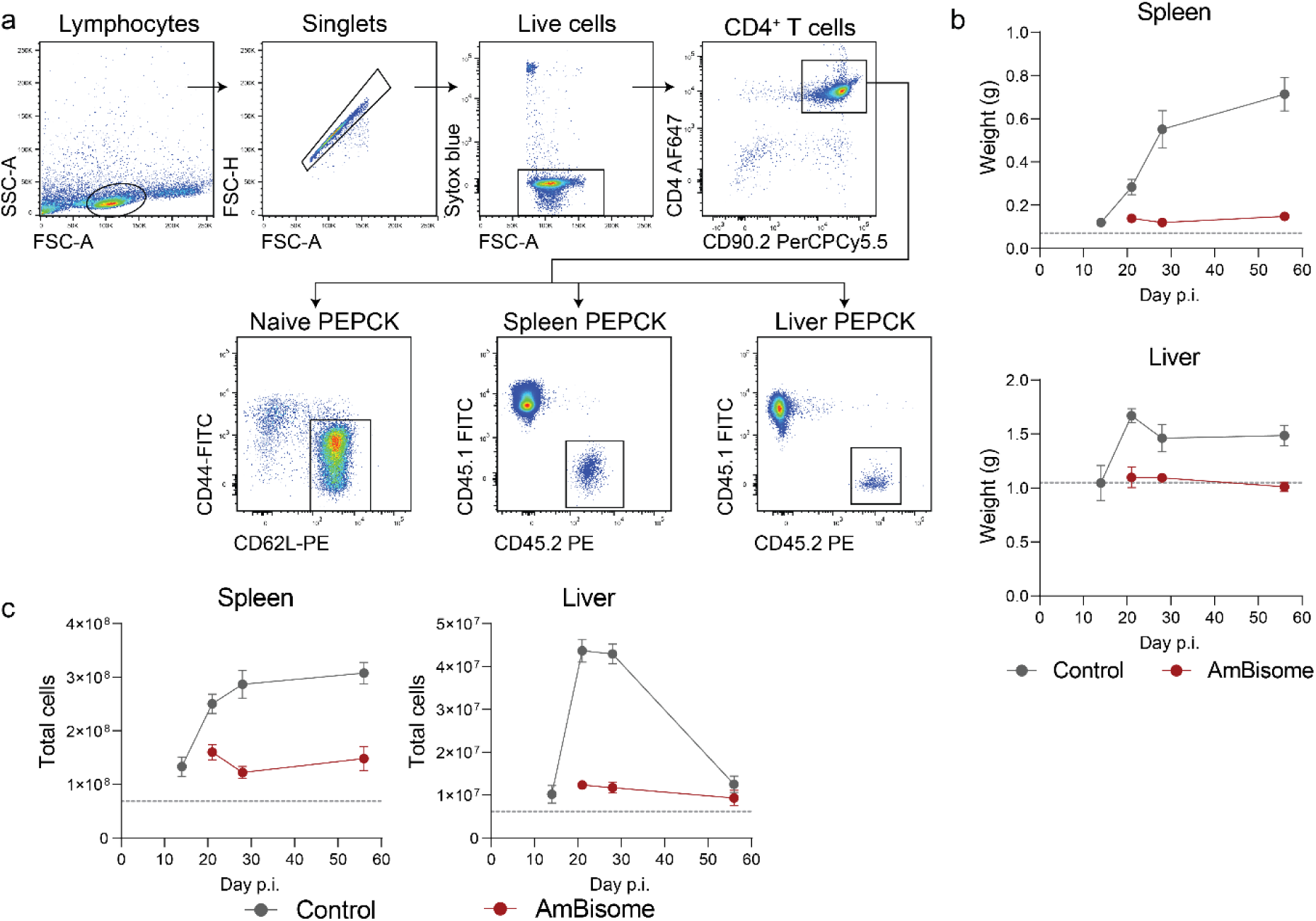
**(a)** Flow cytometry gating strategy for isolating PEPCK cells at day 0 (Naïve PEPCK), or from day 14 p.i. onwards from the spleen and liver (Spleen PEPCK, Liver PEPCK) for scRNAseq assessment. **(b)** Spleen and liver weights and **(c)** the total number of cells per spleen and liver of B6.CD45.1^+^ mice infected with L.donovani treated with vehicle control (control) or AmBisome (n = 3-5 mice per group). Error bars represent mean ± SEM.

**Extended Data Figure 2 (related to Figure 1).**
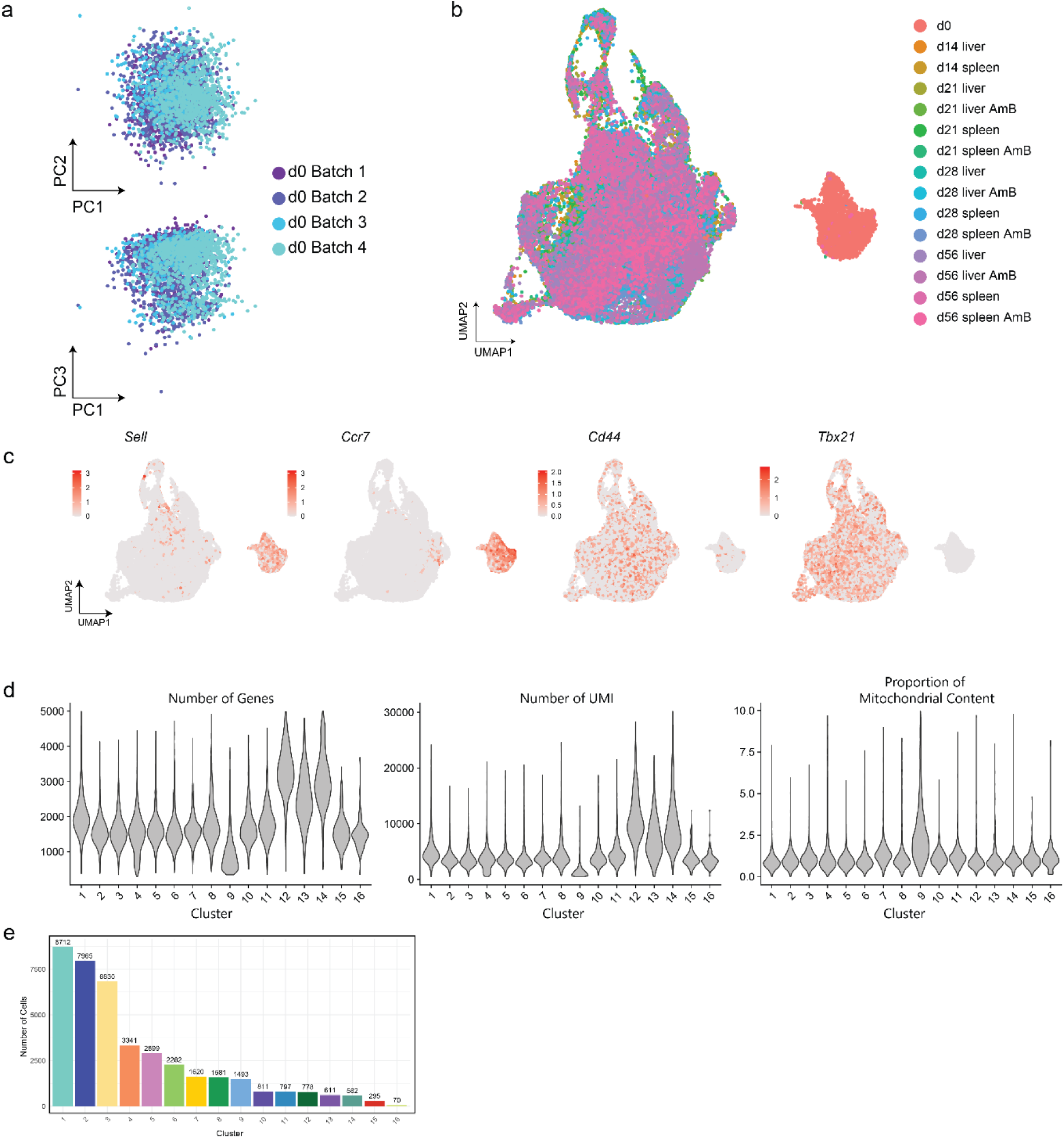
**(a)** Principal component (PC) analysis showing day 0 PEPCK cells from Batch 1-4. **(b)** UMAP visualisation showing PEPCK cells from day 0-56 p.i. from the spleen and liver from vehicle control and AmBisome-(AmB) treated mice. **(c)** UMAP visualisation showing the expression of Sell, Ccr7, Cd44 and Tbx21. **(d)** Distribution of PEPCK cells from day 14-56 p.i. after filtering for number of genes (200<nGenes<5000) and percentage of mitochondrial content (<10%). **(e)** Bar graph showing the number of cells per cluster.

**Extended Data Figure 3 (related to Figure 2).**
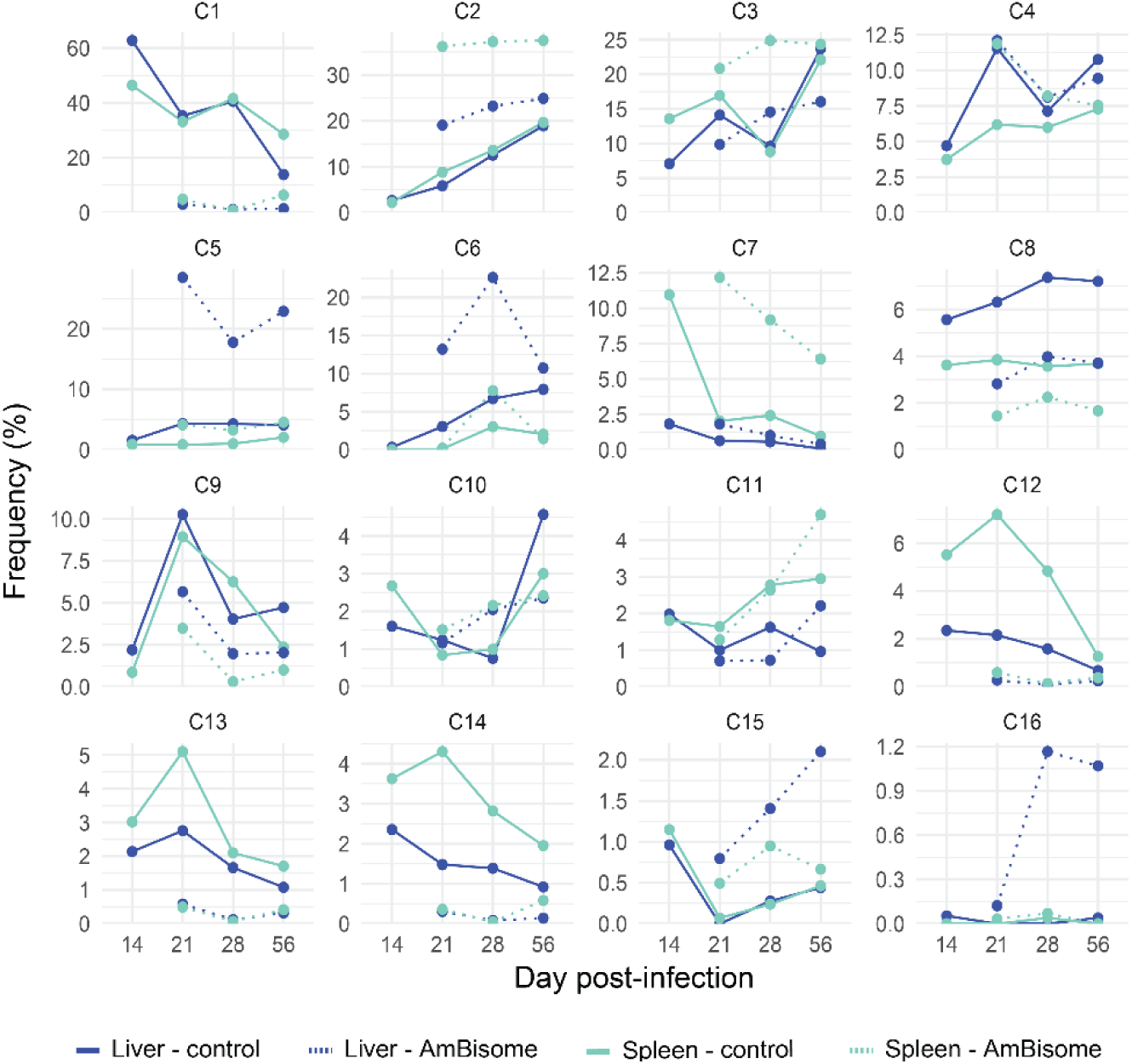
Proportion of total cells per sample for each cluster (C1-C16).

**Extended Data Figure 4 (related to Figure 2).**
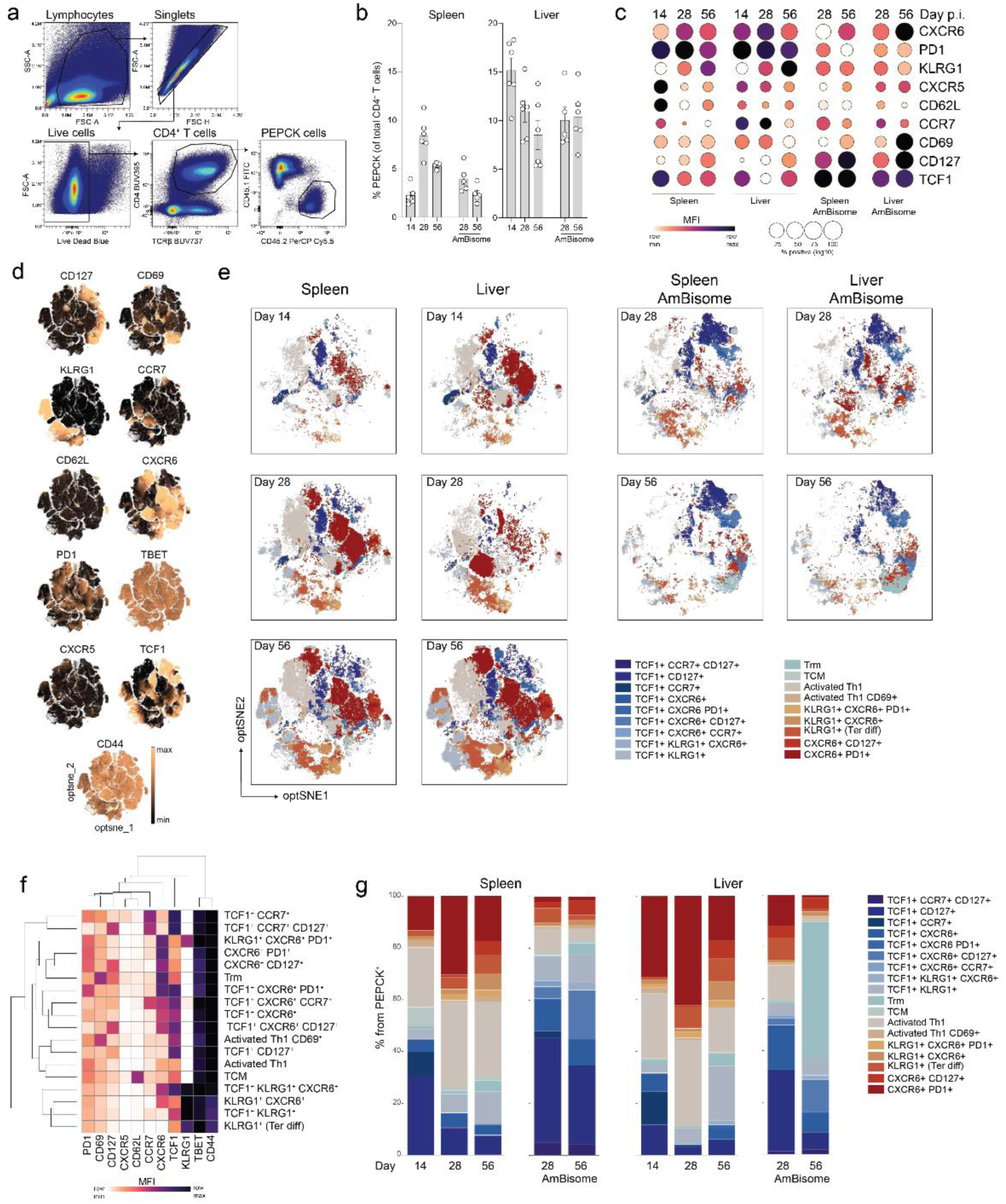
**(a)** Flow cytometry gating strategy to identify PEPCK cells. **(b)** Graph showing the percentage of PEPCK cells in the spleen and liver from day 14 to 56 p.i.. Data are from one experiment, n = 5/6 mice. Error bars represent mean ± SEM. **(c)** Dot plot summarizing the average expression (as mean fluorescent intensity; MFI) and percentage (as postitive events) for PEPCK cells from day 14-28 p.i.. **(d)** optSNE plots showing the expression of CD127, CD69, KLRG1, CCR7, CD62L, CXCR6, PD1, TBET, CXCR5, TCF1, and CD44 on PEPCK cells from day 14 – 56 p.i. from the spleen and liver, from mice treated with vehicle control or AmBisome. **(e)** Clustering of PEPCK cells based on the MFI of all markers shown in (d), **(f)** and the individual molecule expression for each cluster is shown in the heatmap. **(g)** Bar graphs showing the distribution of clusters from (e) for each time point and tissue.

**Extendend Data Figure 5 (related to Figure 3).**
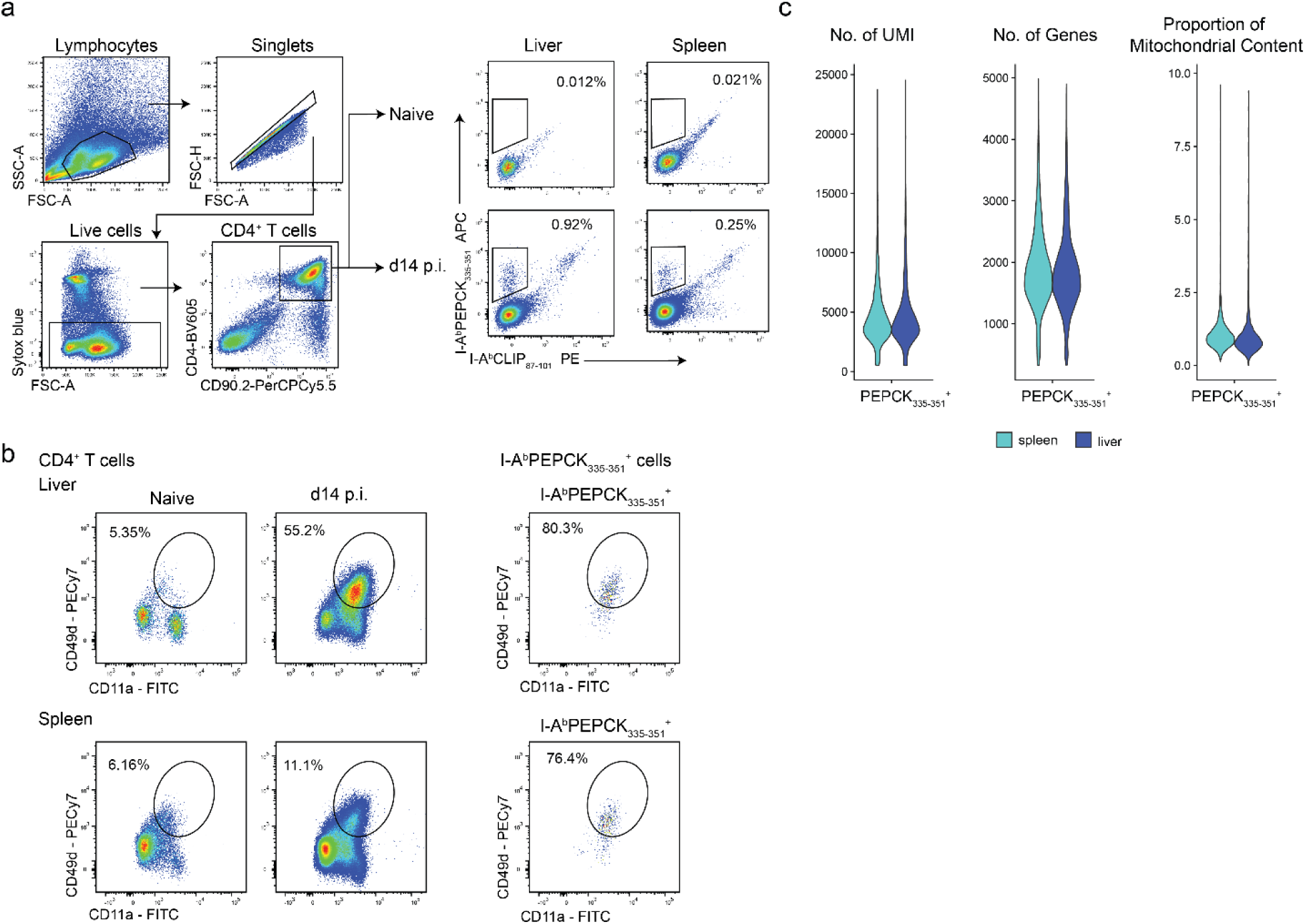
**(a)** Flow cytometry gating strategy used to isolate I-A^b^ PEPCK_335-351_^+^ cells. **(b)** Representative flow cytometry plots showing the expression of CD49d and CD11a on CD4+ T cells (left) and I-A^b^ PEPCK_335-351_^+^ cells (right) in the spleen and liver at day 14 p.i. in experimental VL. **(c)** Violin plots of I-A^b^ PEPCK_335-351_^+^ cells from day 14 p.i. showing quality control metrics.

**Extended Data Figure 6 (related to Figure 4).**
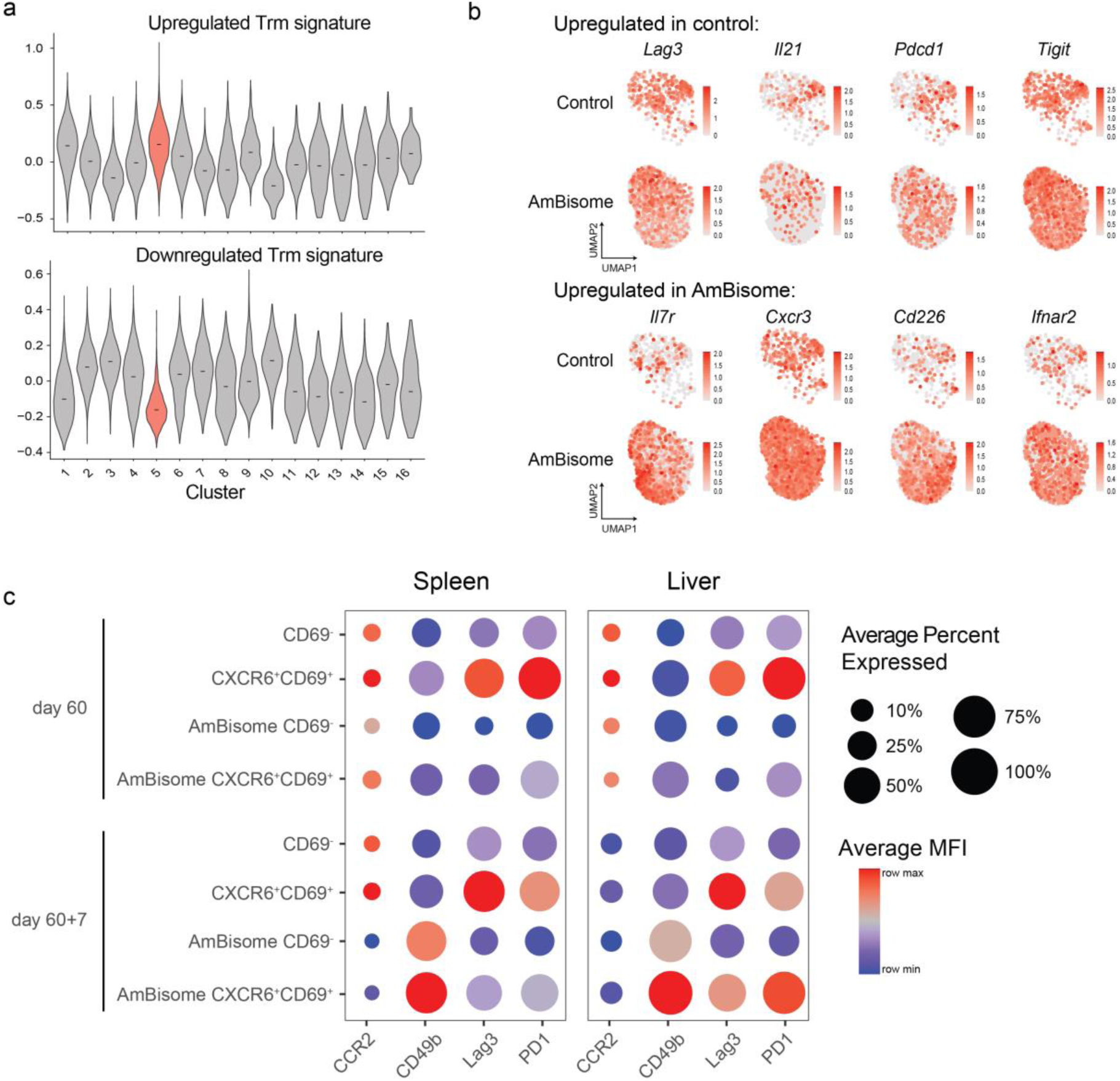
**(a)** Violin plots showing the expression of a Trm gene signature^28^, showing genes that are up- or down-regulated across PEPCK cell clusters. **(b)** UMAP visualisations showing the expression of treatment-associated marker genes in Trm-like PEPCK cells, subsetted by control and AmBisome treatment groups. **(c)** Dotplot showing the average expression (as mean fluorescent intensity; MFI) and average percent expressed (as positive events) of CCR2, CD49b, Lag3, and PD1 on PEPCK cells from the spleen and liver at day 60 and seven days post-rechallenge (day 60+7) in experimental VL. Average MFI scaled for each marker. Experiment performed twice, representative experiment shown.

**Extended Data Figure 7 (related to Figure 5).**
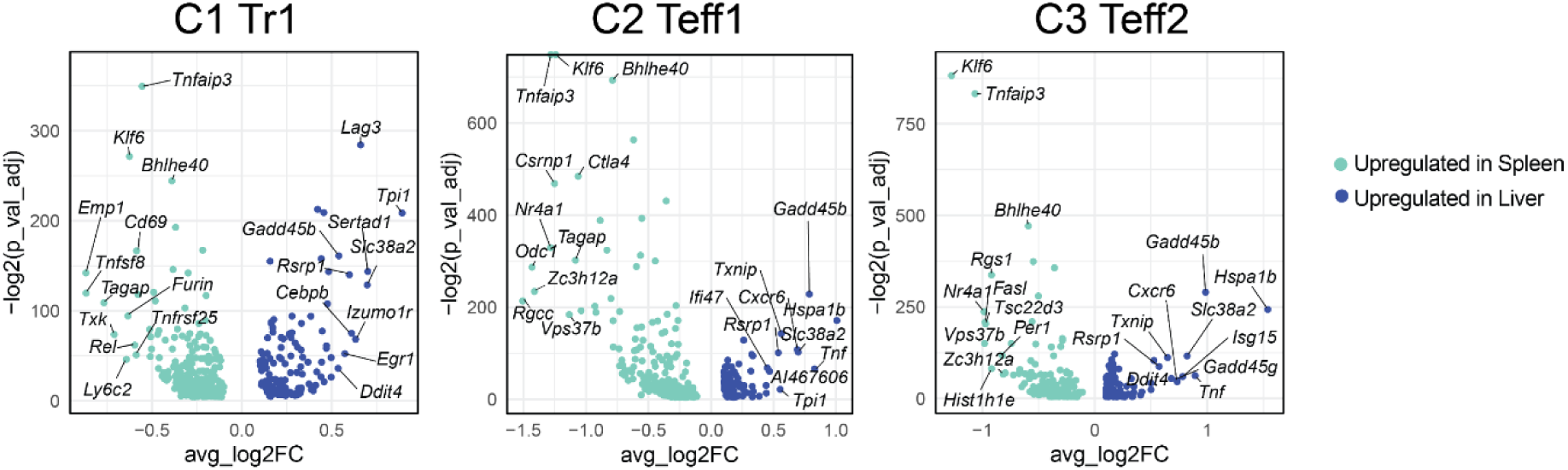
Volcano plots showing log2 adjusted P-values and average log2 fold-change for differentially expressed genes between the spleen and liver PEPCK cells for Cluster 1 (C1 Tr1), Cluster 2 (C2 Teff) and Cluster 3 (C3 Teff2). Each cluster was subsetted and differential gene expression analysis was performed between PEPCK cells from the spleen and liver within each cluster using the “FindMarker()” function in Seurat.

**Extended Data Figure 8 (related to Figure 5).**
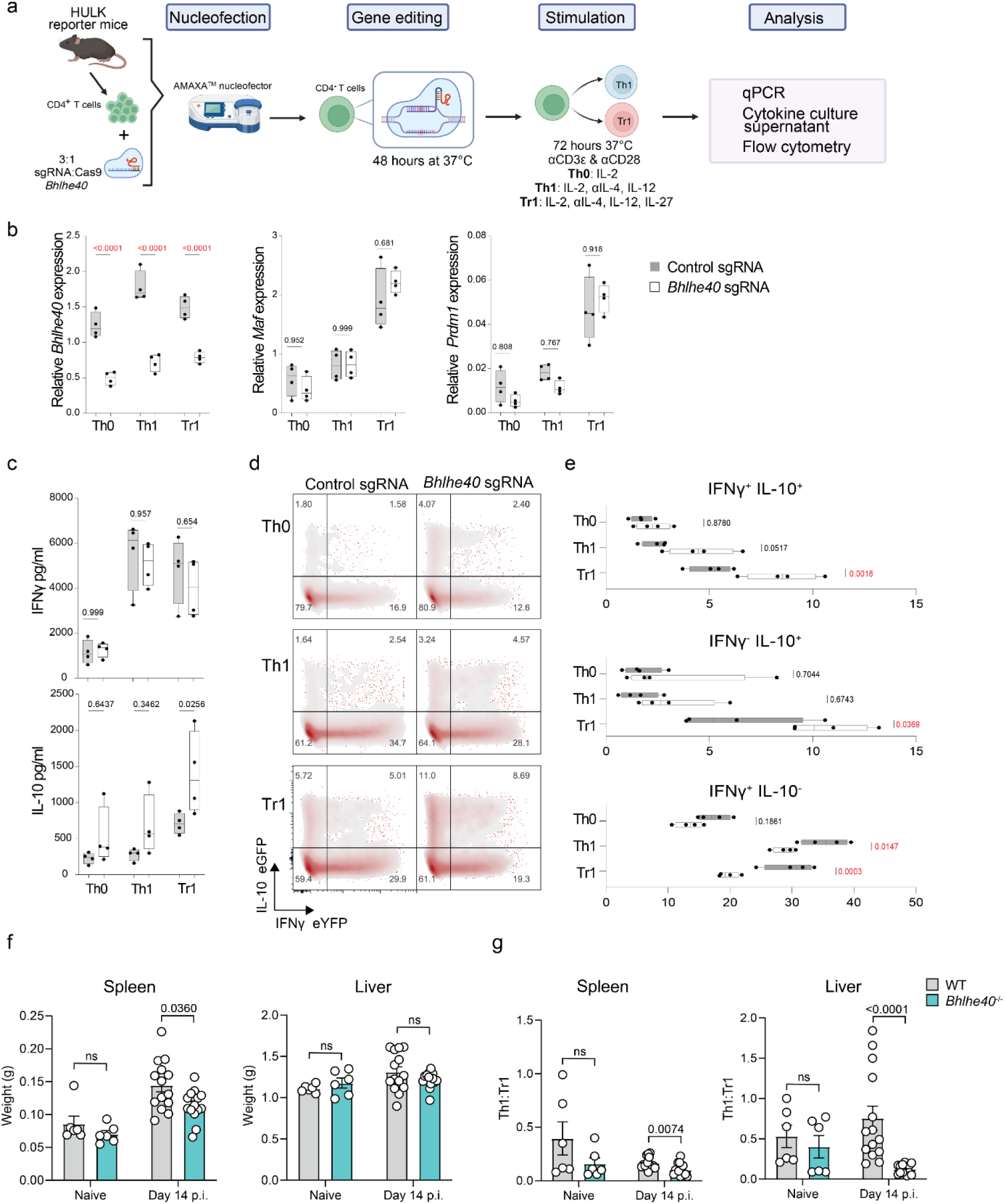
**(a)** CRISPR/Cas9 editing of Bhlhe40 was performed in primary mouse CD4^+^ T cells from triple reporter (Il10gfp x Ifngyfp x Foxp3rfp) C57BL/6 mice, followed by activation with Th0, Th1, Tr1 conditions and assessment. **(b)** Bhlhe40, Maf and Prdm1 mRNA expression by CD4^+^ T cells was detected by qPCR and normalised to the housekeeping gene Hprt mRNA. Box plots show the extent of lower and upper quartiles plus median, while whiskers indicate minimum and maximum data points. **(c)** IFNƴ and IL-10 were measured in cell culture supernatants after 72 hours of activation. **(d** and **e)** Representative flow cytometry plots and enumeration showing the expression of IFNƴ and IL-10 in CD4^+^ T cells after CRISPR/Cas9 editing and activation. n = 4 mice per condition per experiment. Statistical analysis was performed using two-way ANOVA with Sidak’s multiple comparisons test. **(f)** Spleen and liver weights of wild type (WT) and Bhlhe40^-/-^ mice during L. donovani infection at day 14 p.i.. **(g)** Th1:Tr1 cell ratio determined by flow cytometry. Statistical testing was performed using Mann-Whitney U test. Data are pooled from two experiments, where n = 6 naïve WT and Bhlhe40^-/-^ mice, and n =14 WT and Bhlhe40^-/-^ mice at day 14 p.i..

**Extendend Data Figure 9 (related to Figure 6).**
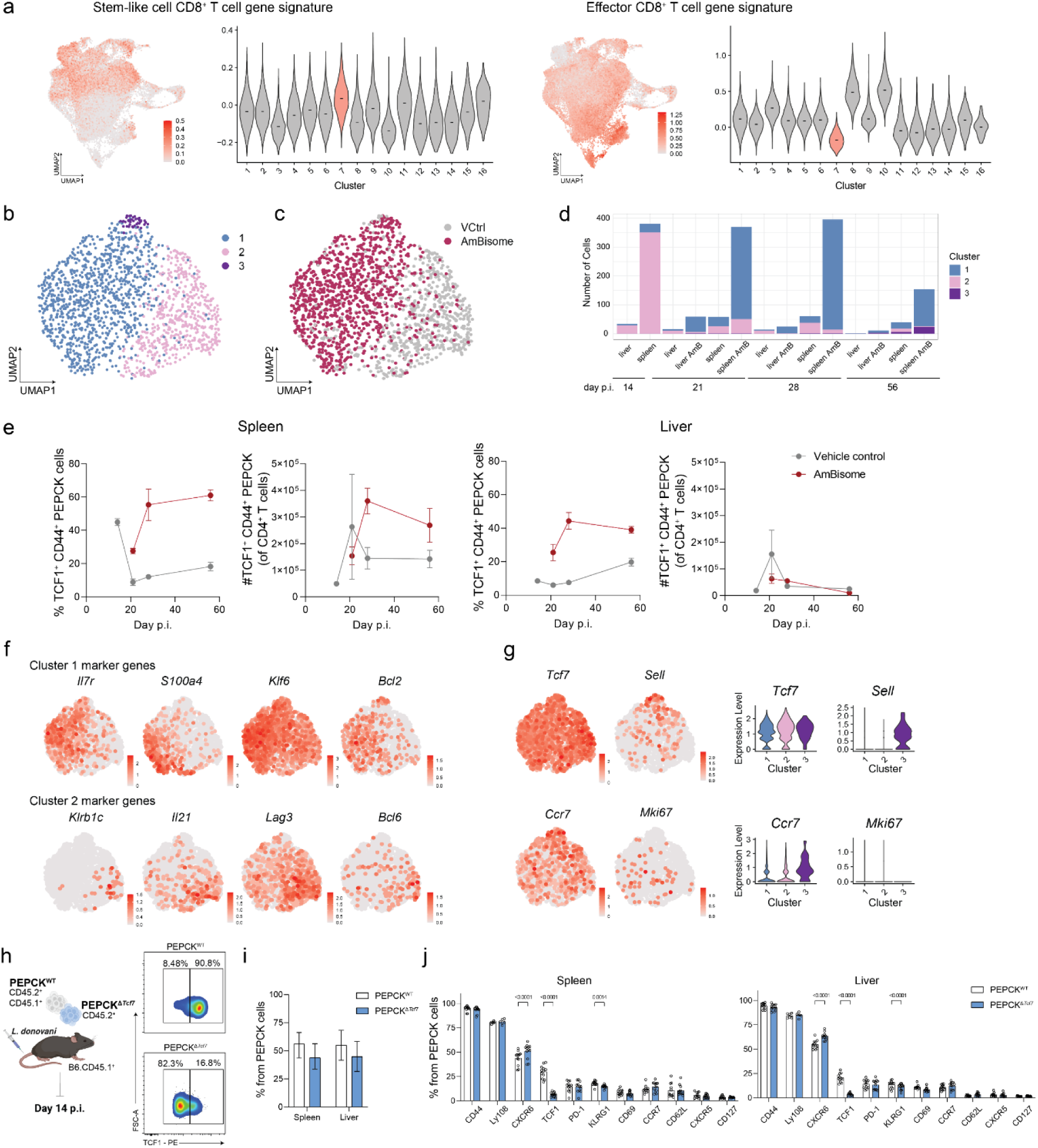
**(a)** UMAP visualisation and violin plot showing the expression of a stem-like and effector CD8^+^ T cell gene signature^56^ on PEPCK cells. **(b)** UMAP visualisation showing unsupervised clustering of stem-like PEPCK cells and **(c)** treatment status. **(d)** Graph showing the number of PEPCK cells in each cluster by sample. **(e)** Graphs showing the percentage and total number of TCF1^+^CD44^+^ PEPCK cells in the spleen and liver during experimental VL. **(f)** UMAP visualisations showing the expression of Cluster 1 and 2 marker genes. **(g)** UMAP visualisations and corresponding violin plots showing the expression of Tcf7, Sell, Ccr7 and Mki67. **(h)** CRISPR/Cas9 editing of PEPCK cells targeting Tcf7 (PEPCK^ΔTcf7^) or a non-specific guide RNA (PEPCK^WT^) was performed and cells were co-transferred into B6.CD45.1^+^ mice, which were then infected with L. donovani. Flow cytometry assessment of PEPCK cells was performed at day 14 p.i.. Flow cytometry plots show the expression of TCF1 on PEPCK^WT^ and PEPCK^ΔTcf7^ cells prior to adoptive transfer. **(i)** Graph showing the frequency of PEPCK^WT^ and PEPCK^ΔTcf7^ cells at day 14 p.i. from the spleen and liver. **(j)** Graphs showing the percentage of surface markers and TCF1 expression on PEPCK^WT^ and PEPCK^ΔTcf7^ cells at day 14 p.i. from the spleen and liver. Data are pooled from two independent experiments, with n = 6 mice per group per experiment. Statistical testing was performed using multiple Mann-Whitney U tests.

**Extended Data Figure 10 (related to Figure 6).**
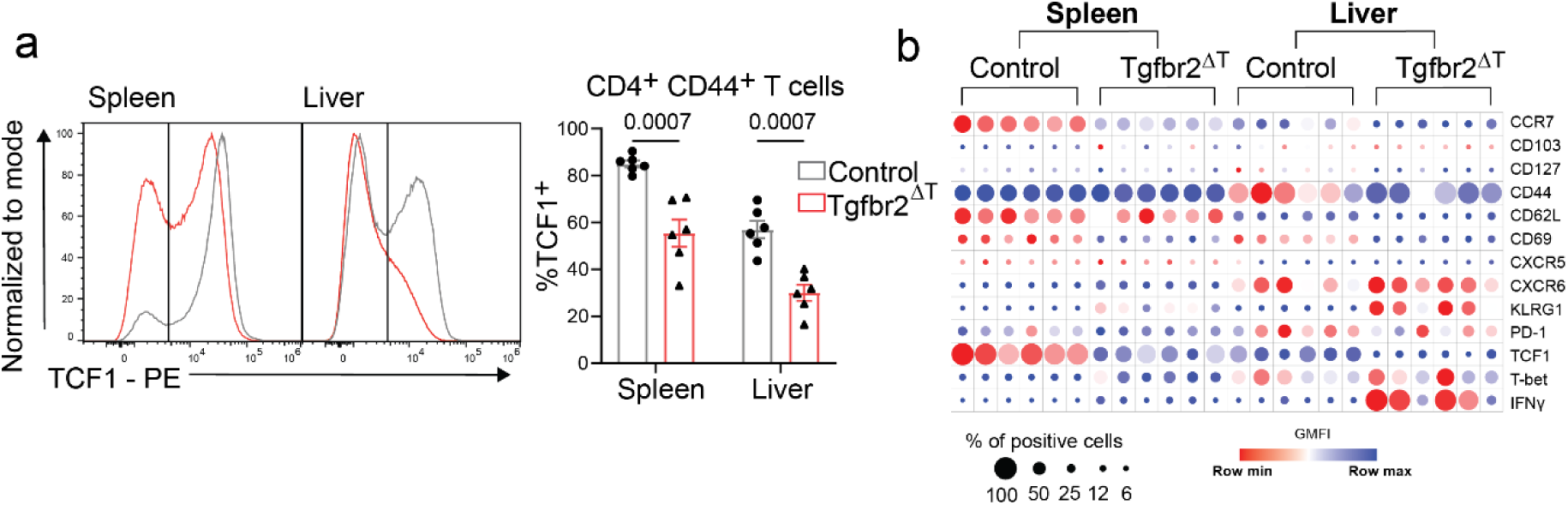
TGFb signalling is needed for efficient development of stem-like CD4^+^ T cells in spleen and liver at day 14 following Leishmania donovani infection. **(a)** CD4^+^ T cells were isolated from the spleen and liver of littermate control (gray lines and columns) and Tgfbr2^ΔT^ (red lines and columns) mice and intracellular levels of TCF1 were measured by flow cytometry. The left panel shows CD4^+^ T cell TCF1 expression and the right hand panel shows frequency of TCF1-positive CD4^+^ T cells in the spleen and liver, as indicated. Statistical testing was performed using a two-tailed Mann-Whitney U test. Error bars represent mean ± SEM. Experiment performed twice, representative experiment shown (n=6 mice per group). **(b)** The average expression (geometric MFI) and frequency (positive events) for listed markers (right-hand side) is summarised in dot-plot format for the spleen and liver, as indicated.

**Extended Data Figure 11. (related to Figure 6).**
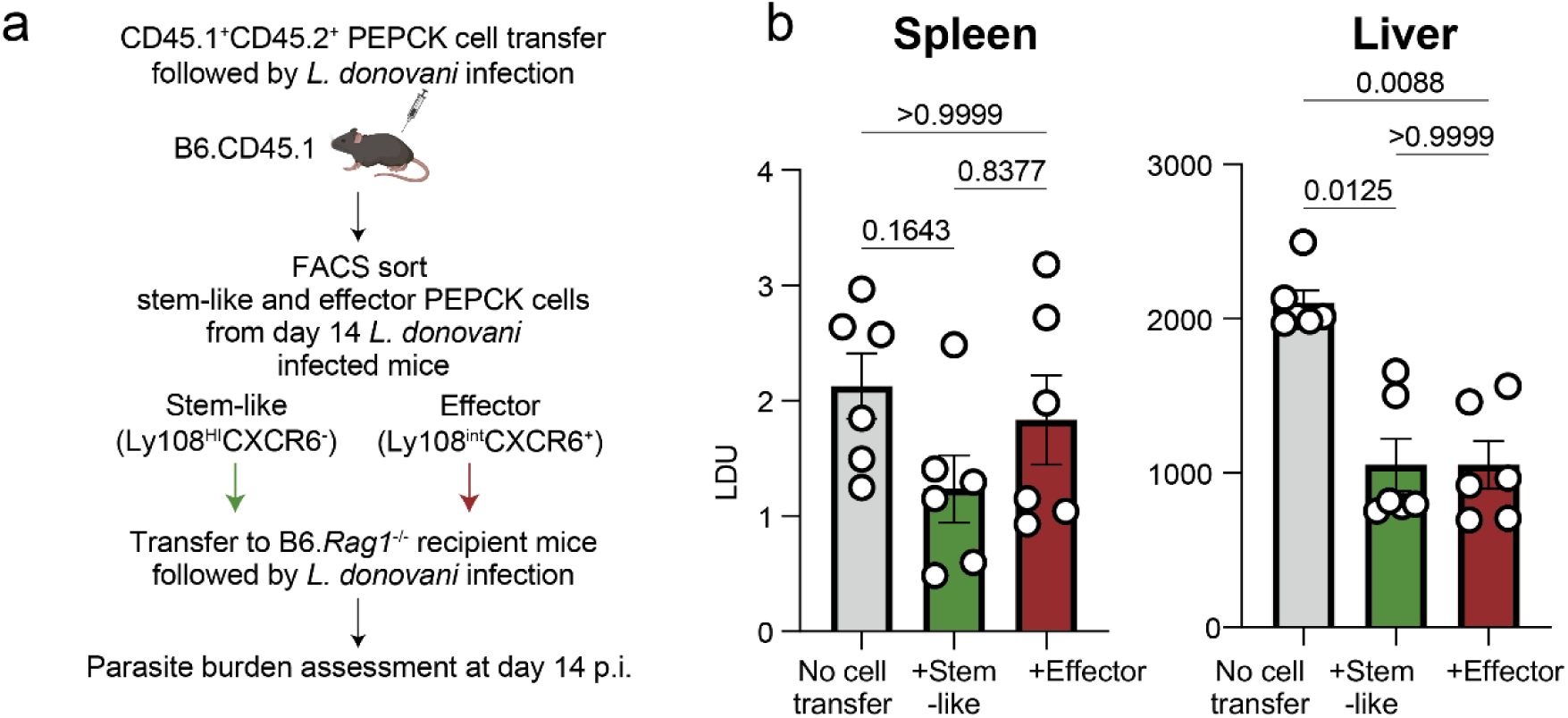
**(a)** Stem-like (Ly108^hi^CXCR6^-^) and effector (Ly108^int^CXCR6^+^) PEPCK cells were isolated from the spleens of L. donovani infected mice at day 14 p.i. and adoptively transferred seperately into B6.Rag1^-/-^ mice, that were infected with L. donovani the day after. A group of B6.Rag1^-/-^ mice were infected but did not receive any transferred cells (No cell transfer), serving as a parasitemia control. **(b)** Parasite burdens of B6.Rag1^-/-^ mice at day 14 post-L. donovani challenge in the spleen and liver. LDU, Leishman-Donovan units. Experiment performed twice, representative experiment shown (n=6 mice per group). Error bars represent mean ± SEM. Statistical analysis was performed using a Kruskal-Wallis with Dunn’s multiple comparisons test.

**Supplementary Table 1:**
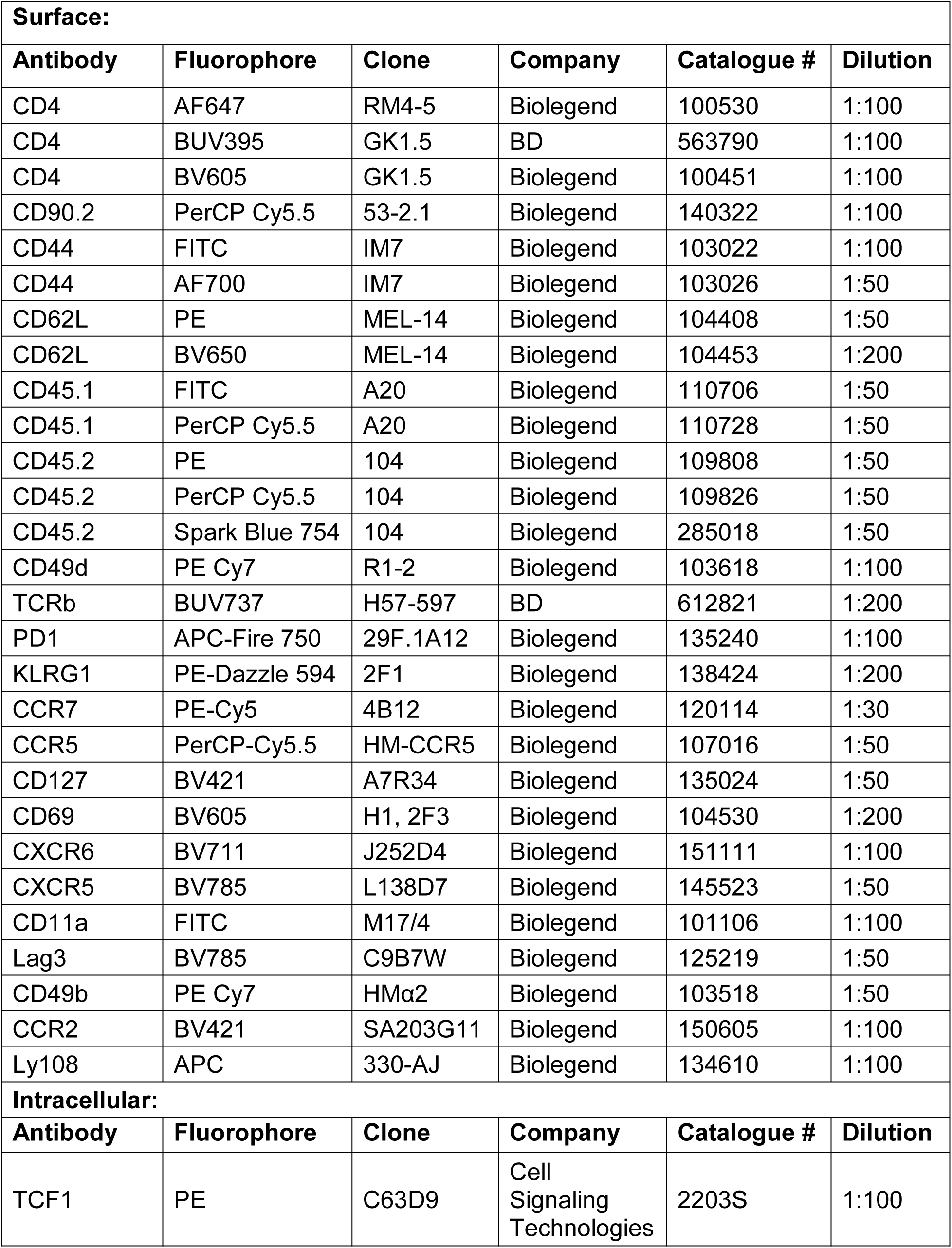
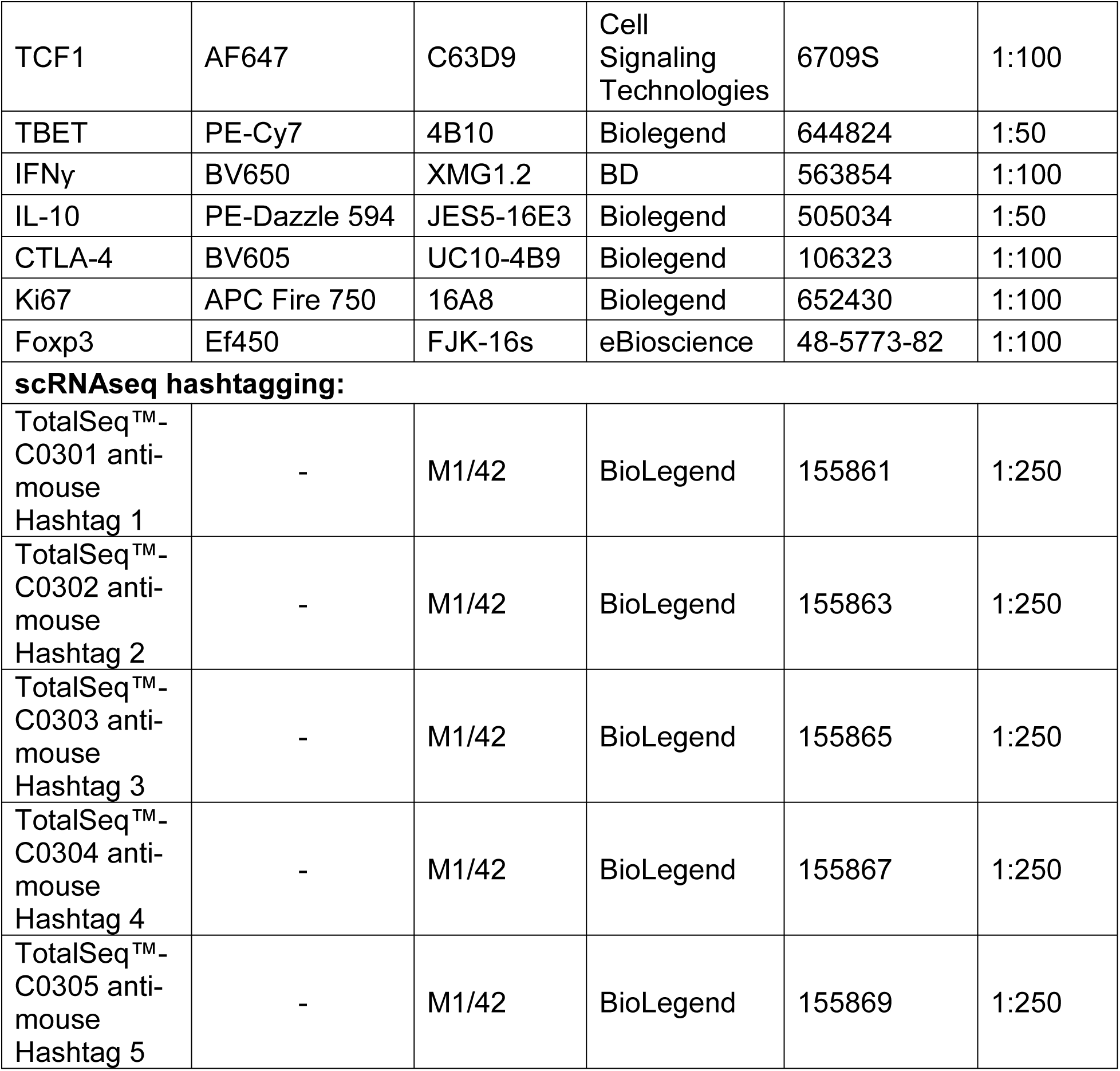
Details of antibodies used.

